# Enhancing protective efficacy and immunogenicity of hemagglutinin-based influenza vaccine utilizing adjuvants developed by BECC

**DOI:** 10.1101/2025.06.03.657703

**Authors:** Sayan Das, Brandon M Tenaglia, Devon Riley, Sydney Speed, Lauren Baracco, Francesca M Gardner, David J Varisco, Hyojik Yang, Haye Nijhuis, Lucas Kerstetter, Florian Krammer, Weina Sun, Lynda Coughlan, Matthew B Frieman, Robert K Ernst

## Abstract

Seasonal influenza viruses continue to pose a significant threat, causing substantial morbidity and mortality in the US and worldwide despite the availability of vaccines and antivirals. These challenges may be addressed by improving vaccine immunogenicity through the inclusion of adjuvants that enhance immune responses against key antigens including influenza hemagglutinin (HA). BECC (Bacterial Enzymatic Combinatorial Chemistry) adjuvants are novel Toll-like Receptor 4 (TLR4) ligands created by modifying enzymes from lipid A synthesis pathways in Gram-negative bacteria. This study compares the ability of the biological and synthetic versions of these adjuvants to enhance the efficacy of recombinant HA (rHA) antigens in mouse influenza virus challenge. Mice immunized with rHA adjuvanted with BECCs stimulate the humoral and cell-mediated arms of the immune system without exhibiting cytotoxicity/pyrogenicity. A robust HA-specific immunoglobulin subtype, especially IgG2a, response was observed in mice adjuvanted with BECCs as compared to control adjuvants, MPL, and PHAD Further, animals adjuvanted with BECC470 cleared infection seven days post-infection, demonstrating their potential for further translational development. Vaccination adjuvanted with BECCs were also able to increase immune recognition of linear B and T cell epitopes when compared to control adjuvants, as well as induce durable immune response eighteen months post-vaccination. Together, these findings indicate that BECCs may serve as highly effective adjuvants in influenza vaccination.

**One Sentence Summary:** Efficacy of engineered lipid A molecules as adjuvants for HA-based influenza vaccine

## Introduction

Influenza A virus (IAV) is a highly contagious respiratory virus resulting in symptoms leading to a spectrum of illnesses ranging from mild to severe, and in some cases, fatal complications. The World Health Organization estimates that seasonal IAV infections are responsible for 3 to 5 million cases of severe illness annually, resulting in 290,000 to 650,000 respiratory deaths worldwide ^1^. In the US, the Centers for Disease Control and Prevention (CDC) reports that influenza has caused between 9.3 million and 41 million illnesses, 100,000 to 710,000 hospitalizations, and 4,900 to 51,000 deaths annually from 2010 to 2023. Overall, while children and younger adults are more likely to become infected, older adults (aged 65 and above) face the highest risk of severe outcomes from seasonal influenza and comprise 50 to 70% of influenza-related deaths^1^. Therefore, prevention through seasonal vaccination remains the focus of combating the global impact of influenza on society.

Influenza A and B viruses are responsible for seasonal influenza epidemics, causing significant morbidity and mortality worldwide each year. While both types contribute to seasonal outbreaks, IAVs have a unique ability to undergo major antigenic shifts, potentially leading to pandemics. The only observed influenza pandemic in the 21st century was caused by a novel IAV (H1N1) strain in 2009, which emerged from a reassortment of human, swine, and avian influenza viruses, highlighting the pandemic potential of IAV ^2^. Antigenic drift, which is responsible for seasonal influenza strains, is caused by the gradual accumulation of mutations in the surface proteins, hemagglutinin (HA) and neuraminidase (NA) that function for viral entry and release, respectively.

These genetic changes enable the virus to bypass previously acquired immunity, driving the ongoing evolution of influenza viruses and requiring yearly updates to flu vaccines to match current circulating strains, while also serving as key targets for vaccine development efforts. ^3^.

The most widely administered influenza vaccines in the United States are egg- and cell-based unadjuvanted quadrivalent vaccines containing four antigens, typically two influenza A and two influenza B strains. Standard-dose egg-based quadrivalent vaccines contain 15 μg of hemagglutinin (HA) per strain, for a total of 60 μg of HA protein per dose. Cell-based quadrivalent vaccines, such as Flucelvax Quadrivalent (Seqirus, USA) contain 15 μg of rHA (recombinant HA) per strain (60 μg total), and the recombinant quadrivalent vaccine Flublok Quadrivalent (Sanofi Pasteur, France) contains 45 μg of rHA per strain (180 μg total). Influenza vaccines containing adjuvants, such as MF59 (a squalene-based oil-in-water emulsion) have been developed to enhance the immune response in populations with reduced vaccine effectiveness, particularly individuals aged 65 years and older or those who are immunocompromised^4^. Adjuvanted vaccines, including FLUAD (Seqirus, USA), are designed to strengthen, broaden, and lengthen the immune response to influenza vaccination, potentially offering improved protection against seasonal influenza strains compared to standard unadjuvanted vaccines in these vulnerable groups ^5,6^.

Depending on the type of innate responses activated, adjuvants can significantly alter both the quality and quantity of adaptive immune responses, enhancing or modulating immune responses when compared with antigen alone ^7,8^. This modulation can lead to improved antibody production, increased T cell responses, and enhanced immunological memory, potentially resulting in more effective and longer-lasting vaccine-induced protection. Currently approved adjuvants for human vaccines, such as aluminum salts and oil-in-water emulsions, predominantly elicit T helper 2 (Th2) immune responses, which may be insufficient for addressing pathogens that demand robust cellular immunity for effective protection. Aluminum salts, while effective in generating antibody responses are less suitable for inducing strong cell-mediated immunity (CMI) or Th1-type antibodies - IgG2a/c and IgG2b, which are crucial for virus clearance^9^. Similarly, oil-in-water emulsions have not demonstrated consistent, long-lasting protection against intracellular pathogens ^10,11^. In contrast, 3-O-deacyl-4’-monophosphoryl lipid A (MPL) promotes a Th1-biased immune response through TLR4 interaction, stimulating pro-inflammatory cytokines and enhancing cellular immunity effective against intracellular pathogens. This leads to reduced morbidity and mortality in vulnerable populations. GlaxoSmithKline (GSK) developed adjuvant systems AS01 (liposome-based) and AS04 (aluminum-MPL co-formulation) to attempt to balance Th1/Th2 responses, with AS01 typically inducing stronger cellular and humoral immune responses. However, MPL presents challenges including manufacturing complexity, heterogeneity of lipid A congeners, potential increased reactogenicity (especially in AS01 formulations), and proprietary control issues.

To enhance adjuvant development, future vaccine formulations require innovative adjuvants or novel formulations that stimulate specific components of the host’s innate immune system. Bacterial Enzymatic Combinatorial Chemistry (BECC) has been employed to create biologically-derived lipid A-based TLR4 agonists, named BECC438b and BECC470b (designated with “b” suffix). These biological preparations demonstrated the ability to enhance both antibody production and cellular immunity when combined with rHA antigens in adult and aged murine challenge models. Unlike MPL PHAD, and alum, BECC-based agonists stimulate a balanced Th1/Th2 immune response, broaden the immune response against conserved rHA stalk domain, and exhibit significant dose-sparing effects ^8^. Specifically, when formulated with biologically-derived BECCs, a 7-fold reduction in rHA antigen (40 ng) was protective against IAV challenge, as evidenced by mitigation of weight loss, viral titer reduction, and reduced adverse lung pathology. Moreover, a 10-fold decrease in adjuvant dosage still afforded superior protection. Additionally, complete protection was achieved with a prime-only vaccination schedule, meeting the criteria for an effective seasonal vaccine ^8,12^.

To advance the development of the BECC adjuvant platform, synthetically-derived structures of the lead BECC molecules BECC438s and BECC470s (designated with “s” suffix) were generated. The immunogenicity of these synthetic versions were compared to their biologically-derived counterparts and control adjuvants MPL (biological, Invivogen) and PHAD (synthetic, Croda) in a murine challenge study using an influenza rHA (H1) antigen. The comparison between biologically-derived and synthetically-derived adjuvants was crucial due to their distinct molecular compositions. While biologically-derived preparations contain multiple congeners, offering a diverse range of molecular structures, synthetically-derived molecules have an identical acyl chain composition but differ in their levels of phosphorylation. This approach allows for a more controlled evaluation of the adjuvants’ efficacy and potentially addresses issues related to consistency and manufacturing complexity.

In this study, BECC adjuvants (biological or synthetic) demonstrated enhanced efficacy compared to the control adjuvants, with BECC470b and BECC470s exhibiting the highest protective efficacy against influenza (H1) in a murine airway challenge model. These formulations induced robust humoral responses, evidenced by increased HA-specific IgG1 and IgG2a serum antibodies, as well as strong cell-mediated immune responses, as shown by cytokine secretion following *ex vivo* stimulation of splenocytes with rHA and *in vivo* viral infection. Notably, both BECC470b and BECC470s elicited long lasting immune responses. These findings underscore the potential of synthetically-derived BECC molecules as promising candidates for next-generation vaccine adjuvants, offering improved efficacy and potentially addressing manufacturing and consistency challenges associated with biologically-derived adjuvants.

## Results

### Structures of synthetically-derived BECCs

Biologically-derived BECC438 and BECC470 have been previously identified and characterized as potent adjuvants ^13–15^. However, to address the issue of batch-to-batch variations resulting from a mixture of congeners, a single structure for each molecule was selected and chemically synthesized. As shown in **Figure 1**, synthetic BECC438 and BECC470 are hexa-acylated structures with a symmetrical (3 + 3)-type acylation pattern consisting of four 3-OH C14 acyl chains attached to the diglucosamine backbone and a secondary C16 -chain added at the 2 position and a C16:1 acyl chain added at the 3’ position. The key distinction between BECC438s and BECC470s lies in their phosphorylation states. BECC438s contains two phosphate groups at the 1 and 4’ positions, whereas BECC470s is mono-phosphorylated with a phosphate group only at the 1’ position. The structures of the synthetically-derived molecules were confirmed using tandem-MS analysis (**Fig. S1**), ensuring their conformity to the desired specifications and providing a more controlled basis for evaluating their efficacy as vaccine adjuvants.

**Fig. 1.**
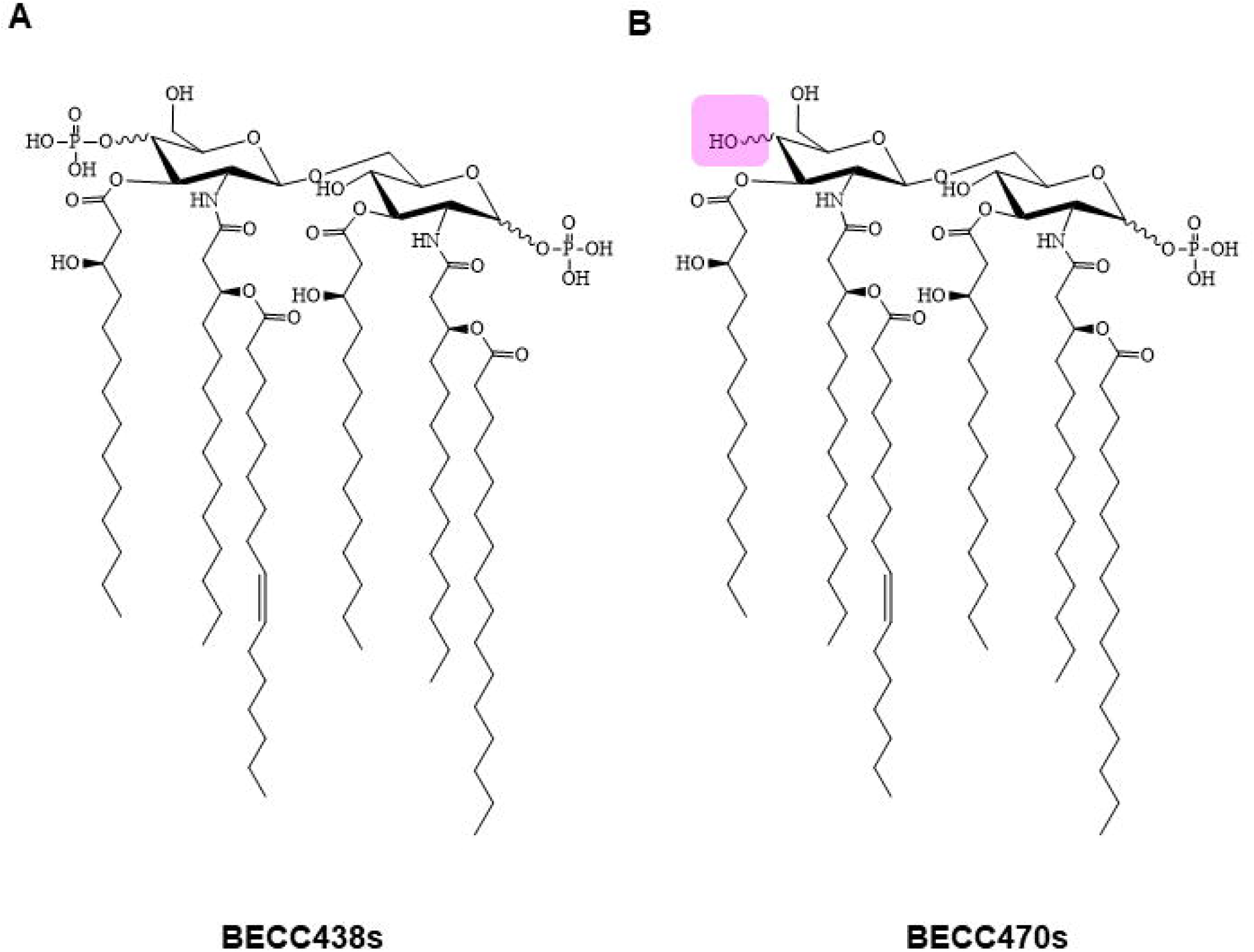
Structures synthetically-derived BECCs. Chemical structures of BECC438s (**A**) and BECC470s (**B**). Pink box marks the location of the missing 4’ phosphate group of BECC470s.

## BECC adjuvanted rHA induces robust serum IgG response in immunized mice

To evaluate the adjuvant potential of biologically- and synthetically-derived BECC438 and BECC470, mice were contralaterally immunized on hind limbs with 40 ng of rHA formulated (admixed) with 50 µg of PHAD, MPL, BECC438b, BECC438s, BECC470b, and BECC470s in a prime (day 0) - boost (day 14) schedule (**Fig. 2A**). Control groups received PBS (sham) or rHA without adjuvant. The 40 ng dose of rHA was selected based on previous studies indicating its non-protective nature when administered alone yet demonstrating the potential for enhanced efficacy when combined with BECC adjuvants ^8^. MPL (biological) and PHAD (synthetic) served as comparators.

**Fig. 2.**
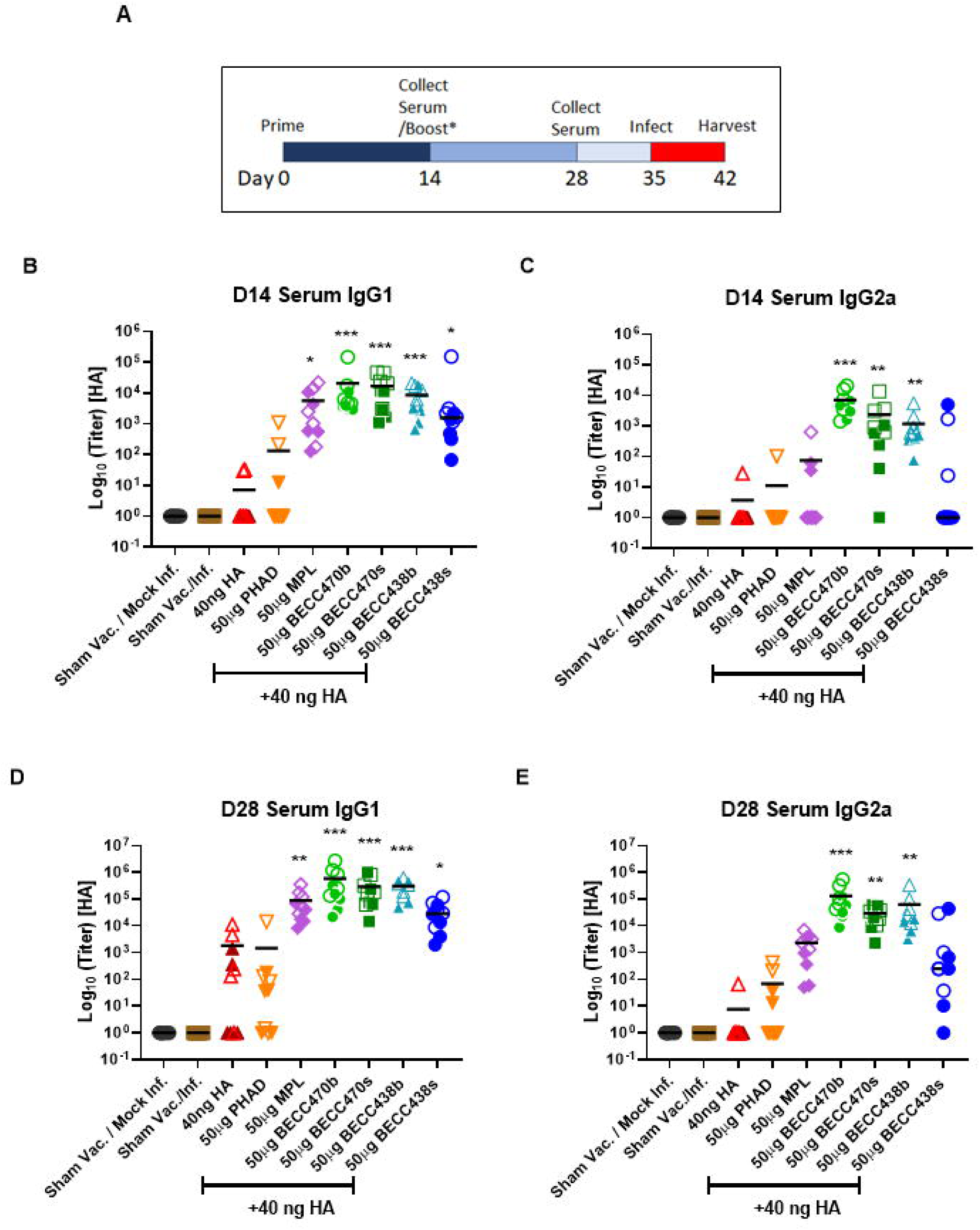
rHA adjuvanted with BECC induce a robust humoral immune response in mice. 6-week-old BALB/c mice were immunized intramuscularly on days 0 and 14 with 40 ng of rHA with indicated adjuvants. Sham groups were immunized with PBS. On days 14 and 28 post-immunization, serum IgG1 (**B and D**) and IgG2a (**C and E**) were assessed by ELISA. Each group consisted of 5 males (solid symbols) and 5 females (open symbol). Data were analyzed by Kruskal-Wallis test by comparing to Sham Vaccinated/infected group. *p<0.05, **p<0.01, ***p<0.001.

To compare Th1 and Th2 immune responses, serum levels of HA-specific IgG1 (Th2) and IgG2a (Th1) were measured on days 14 and 28, revealing significant differences among immunized groups. On day 14, rHA adjuvanted with MPL (p=0.127), BECC470b (p < 0.001), BECC470s (p < 0.001), BECC438b (p < 0.001), and BECC438s (p = 0.036) showed significantly higher HA-specific IgG1 antibody titers compared to sham groups (**Fig. 2B**). In contrast, only BECC470b (p < 0.001), BECC470s (p = 0.004), and BECC438s (p = 0.003) groups showed statistically significant IgG2a levels (**Fig. 2C**). By day 28, all BECC-adjuvanted groups demonstrated a 1.5-2 log increase in HA-specific IgG1 and IgG2a titers compared to day 14 levels (**Fig. 2D-2E**). Among the control adjuvants, only MPL-induced IgG1 was significantly higher than PBS-immunized groups. These results indicate that BECC adjuvants generate a robust humoral immune response 14 days post-prime, which was further enhanced post-boost at day 28. Notably, no significant differences were observed in antibody responses between male (solid symbols) and female (open symbols) mice (**Fig. 2B-E**).

## Protective efficacy of rHA adjuvanted with BECCs against H1N1 challenge (NL09)

Mice vaccinated with different HA-adjuvant formulations were challenged with NL09 (H1N1) to assess the correlation between antibody levels and protection. Following intranasal challenge with 3,200 PFU of NL09 (H1N1), BECC470b-vaccinated mice maintained weight profiles comparable to uninfected controls, confirming robust protective efficacy, whereas BECC470s-, BECC438b-, and MPL-adjuvanted groups showed delayed recovery (approximately day 6 post-challenge), with MPL displaying more severe initial weight loss prior to rebound (**Fig. 3A**). In contrast, the PHAD-adjuvanted and unadjuvanted rHA groups exhibited progressive weight loss without recovery. In contrast, PHAD-adjuvanted and unadjuvanted rHA groups displayed a continuous downward trend in body weight. These results highlight the efficacy of BECC adjuvants, particularly BECC470b/s, against weight loss in this homologous challenge model, outperforming both control adjuvants and unadjuvanted HA.

**Fig. 3.**
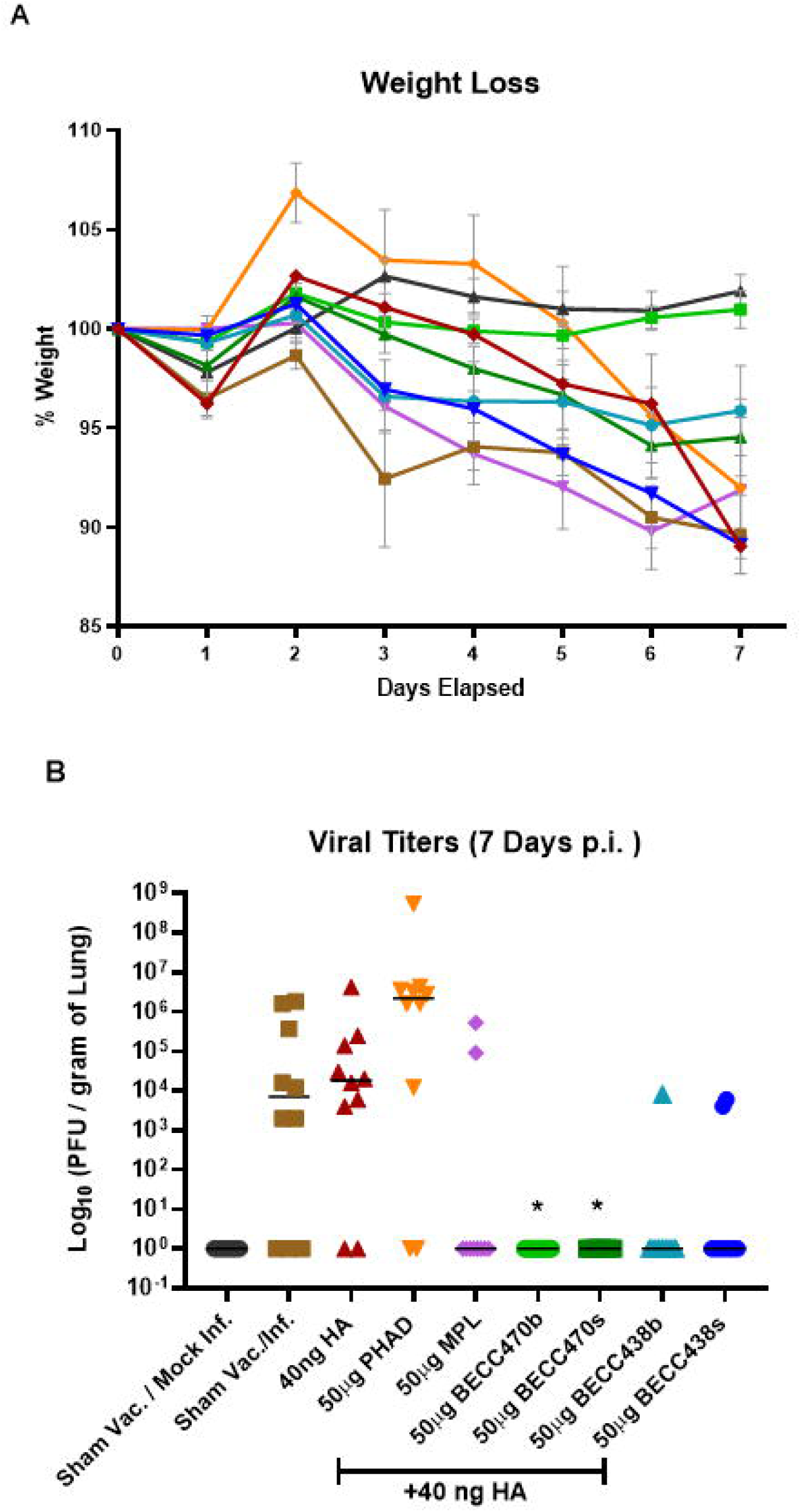
rHA adjuvanted with BECC protected mice from influenza (NL09) challenge. Mice from the previous experiment were intranasally challenged with 3200 PFU of NL09 (H1N1) on day 35 post-immunization. On day 7 post-infection (p.i.), mice were sacrificed and viral titers in lungs were determined (**A**). Body weights were monitored for the infected mice (**B**). Data were analyzed by Kruskal-Wallis test by comparing to Sham Vaccinated/infected group. *p<0.05, **p<0.01, ***p<0.001.

Seven days post-infection, mice were euthanized, and lung viral titers were determined by plaque assay, revealing varying degrees of efficacy in viral clearance among the individual adjuvants. BECC470, in both its biological and synthetic forms, demonstrated a significant reduction in viral loads compared to sham vaccinated/infected groups, with mice adjuvanted with BECC470b and BECC470s exhibiting complete viral clearance and no detectable viral titers in their lungs (Fig. 3B). BECC438b demonstrated high efficacy with only one mouse exhibiting detectable viral titers, while BECC438s and MPL showed moderately reduced effectiveness, and both PHAD-adjuvanted and unadjuvanted rHA groups demonstrated markedly lower effectiveness in clearing the viral infection with eight mice in each group still harboring detectable viral titers. These results correlated with the higher levels of anti-HA IgG1 and IgG2a observed in BECC-adjuvanted groups compared to other groups (**Fig. 2**). The findings collectively demonstrate that BECC-adjuvanted groups elicit a balanced Th1/Th2-type immune response. Consequently, these groups provide superior protection against weight loss and virus replication when challenged with a homologous viral strain.

## Cytokine induction by BECC adjuvanted rHA pre-challenge and post-challenge

TLR4-mediated inflammation involves key pro-inflammatory cytokines that are essential for initiating and sustaining the immune response, thereby contributing to the control of viral loads during infection. The observation that MPL and BECC438b produced high antibody titers but failed to eliminate virus in lungs highlights the importance of investigating the cell-mediated immune responses elicited by each adjuvant. To comprehensively assess the immune response, *ex vivo* cytokine profiles were determined in separate experiments, both pre-challenge and post-challenge. The pre-challenge measurements aimed to evaluate the total immune response generated by the vaccine and adjuvant combination, while the post-challenge measurements focused on assessing the immune response specifically triggered by the viral infection. This approach allows for a more nuanced understanding of how different adjuvants shape both the initial immune response to vaccination and the subsequent response to viral challenge, providing valuable insights into their mechanisms of action and overall efficacy in enhancing vaccine-induced protection.

To assess pre-challenge cytokine responses, mice were immunized as outlined (**Fig. 2A)** and euthanized on day 35 post-immunization. Spleens and lungs were harvested, and single-cell suspensions were prepared and stimulated with rHA for 48 hours at 37°C, 5% CO_2_ in a humidified incubator. Supernatants were collected and analyzed for cytokines using a multiplex assay (U-plex; Meso Scale Discovery). Following normalization, **Fig. 4A** demonstrates the induction of key cytokines (TNF-α, IL-17a, IL-12p70, IFN-γ, IL-6, and KC/GRO) in HA-stimulated splenocyte supernatants with actual values presented in **Fig. S2.** Statistically significant levels of IL-17a were observed in groups immunized with BECC470s (p <0.0001), BECC470b (p <0.0001), PHAD (p = 0.0013), and MPL (p = 0.0018). Notably, BECC470b and BECC470s induced higher levels of cytokines as compared to PHAD and MPL (**Fig. S2**); the remaining cytokines exhibited no statistically significant differences. Lung samples stimulated with rHA exhibited low cytokine levels with no discernible pattern between groups (data not shown).

**Fig. 4.**
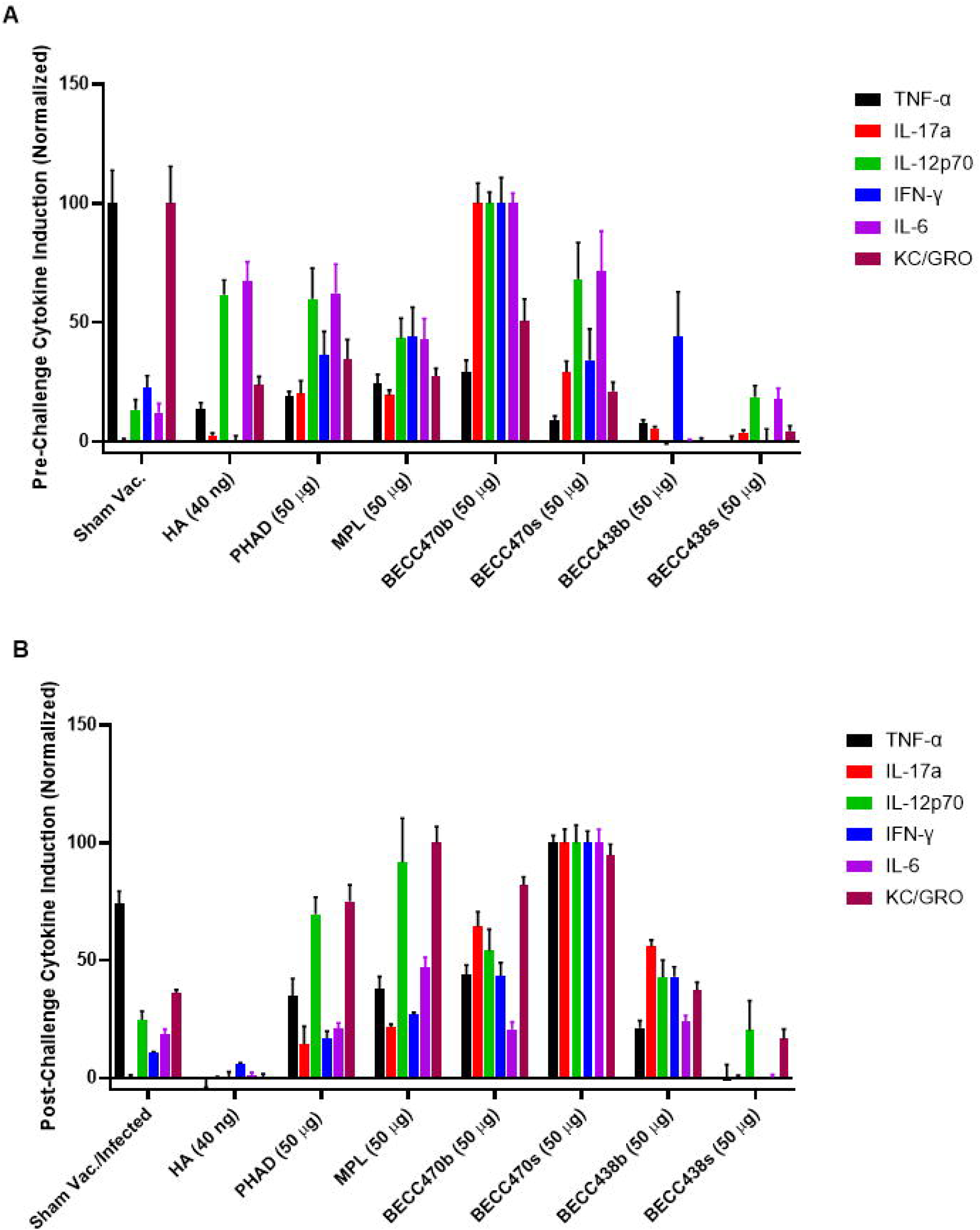
Cytokine induction by BECC adjuvanted rHA pre-challenge and post-challenge. Six-week-old BALB/c mice (n=5/group) were immunized on day 0 and day 14. On day 35 post immunization, mice were sacrificed, and single suspensions of spleen were prepared. Cell concentrations were equalized and were plated onto 96 well plates in the presence of rHA. Supernatants were removed after 48 hours and analyzed for presence of cytokines. The normalized cytokine levels are shown (**A**). A separate group of mice were intranasally challenged with 3200 PFU of NL09 (H1N1). Two days post challenge mice were sacrificed, and single suspensions of lungs were prepared. Cell concentrations were equalized and were plated onto 96 well plates. Supernatants were removed after 48 hours and analyzed for presence of cytokines. **B** shows the normalized cytokine levels.

To investigate the post-challenge cytokine pattern, immunized mice were infected with Influenza A, and their cytokine secretion profile was studied. Mice were vaccinated as previously described and challenged intranasally with 3200 PFU of NL09 (H1N1) on day 35 post-immunization. Two days post-infection, mice were euthanized, and their spleens and lungs were harvested as above. Single-cell suspensions of these organs were prepared, and the supernatants were analyzed for cytokines using a multiplex assay. The normalized cytokine levels are shown (**Fig. 3B)**.

All adjuvanted groups, MPL (p = 0.0012), BECC470b (p <0.0001), BECC470s (p <0.0001), and BECC438b (p <0.0001) showed significantly elevated IL-17a as compared to the sham control. In contrast to pre-infection results, statistically significant levels of IFN-γ were observed in BECC470b (p = 0.0010), BECC470s (p <0.0001), and BECC438b (p = 0.0012) adjuvanted groups (**Fig. S3**). Interestingly, the cytokines detected in the supernatants of post-infection lungs were similar to those detected in pre-infection spleens, suggesting the homing of HA-specific immune cells from spleens to lungs. Moreover, the magnitude of cytokines secreted was higher in post-infection lungs compared to HA-stimulated pre-infection spleens. These results indicate that BECCs may be crucial in inducing Th1/Th17 cytokines, thereby protecting mice from infection.

## Identification of linear B and T cell epitopes

BECC-based adjuvants drive robust immunity, eliciting high titers of HA-specific antibodies (**Fig. 2B–E**). To dissect the humoral (B cell) response, immune serum was screened against a peptide library, 139 overlapping 15-mers (11-aa overlap) providing a detailed map of antibody binding across the HA protein. This epitope profiling revealed adjuvant-specific differences in immunodominant regions, suggesting BECC formulations may broaden epitope targeting compared to traditional adjuvants. Comparative analysis showed adjuvant-driven differences in epitope targeting breadth, suggesting BECCs may enhance response diversity as compared to conventional adjuvants.

Briefly, pools of 10 sequential peptides were created (pools 1-14) and coated onto 96-well plates using -Ethyl-3-(3-dimethylaminopropyl)carbodiimide (EDC) to covalently link the free carboxylic group of the protein to the amino group of the microtiter plates. This approach allowed for a comprehensive analysis of antibody binding across the entire HA protein sequence. Pooled serum from mice vaccinated as above was added to each plate, and peptide specific IgGs (total) were detected with peptides in pools 5, 6, and 7 recognized by serum from immunized mice. To identify immune-dominant epitopes, individual peptides from these pools were coated onto 96-well plates, and IgG1 and IgG2a levels were determined (**Fig. 5A and B**). All adjuvanted groups displayed IgG1 responses, with the strongest IgG2a response observed in BECC470b and BECC470s adjuvanted groups. Using a cut-off O.D. value of three times that of the PBS-vaccinated control group, BECC438b had the highest number of peptides (26) above the threshold for IgG1, followed by BECC470s (24) and BECC470b (16); whereas PHAD and MPL showed only 3 and 5 peptides meeting the criteria, respectively. For IgG2a, BECC470s and BECC470b had the highest number of peptides (28 and 27, respectively) above threshold. Both versions of BECC438 and PHAD had considerably fewer peptides meeting the criteria, while rHA only and MPL groups had none. These observations suggest that BECC adjuvants enhance immune recognition of linear HA epitopes, which are located on the head region of the protein **(Fig. 5C)**.

**Fig. 5.**
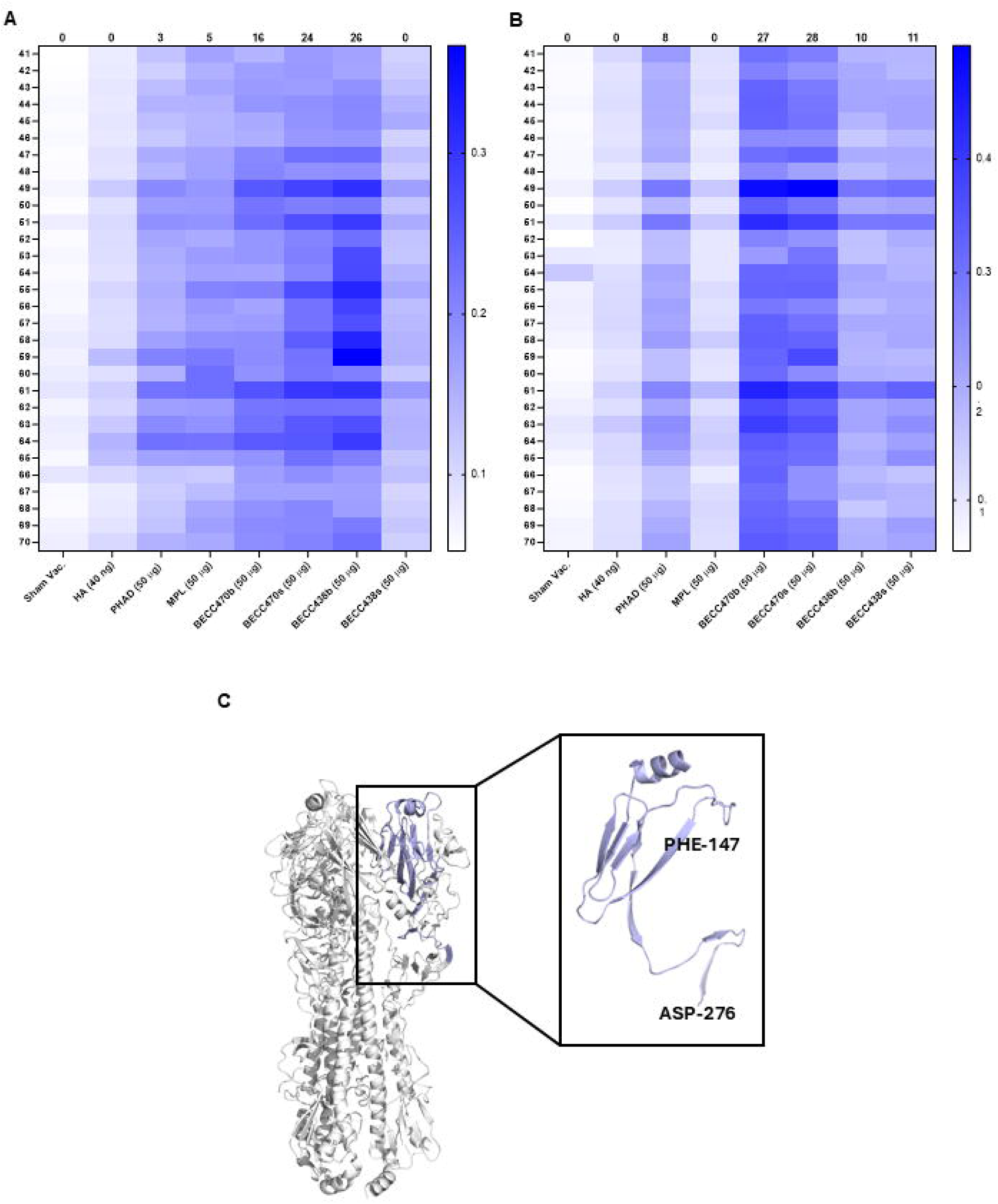
Identification of linear B cell epitopes. Six-week-old BALB/c mice (n=5/group) were immunized intramuscularly on days 0 and 14 with 40 ng of rHA with indicated adjuvants. Sham groups were immunized with PBS. On day 28 post-immunization, mice were bled, serum isolated and IgG1 (**A**) and IgG2a (**B**) titers against individual peptides were determined by ELISA. The heat map shows the O.D. values obtained at 1:50 serum dilution of sera. Numbers on top of each column represents peptides that have O.D. 3 times the respective sham vaccinated control. The location of B cell epitopes on the HA trimer is shown in panel **C**.

To better understand antigen-specific cellular (T cell) immune responses, linear T cell epitope mapping was performed as described above. On day 35 post-immunization, mice were euthanized, spleens harvested, and single-cell suspensions were stimulated with 14 peptide pools. Supernatants were analyzed for IL-17a, a cytokine strongly induced by BECC adjuvants, revealing their potent immunostimulatory effects. (**Fig. 4**). IL-17a was induced in adjuvanted rHA groups, with peptide pools 4 and 6 showing significant induction. Further characterization using 20 individual peptides from these pools revealed that BECC470b and BECC470s elicited the most robust IL-17a response, followed by BECC438s (**Fig. 6A**). Interestingly, peptide 55 located on the head region (**Fig. 6E**), only stimulated the group vaccinated with biologically-derived BECC438b and BECC470b. Control adjuvants showed minimal IL-17a induction without a discernible pattern. This experiment demonstrated that rHA adjuvanted with BECC470b has the most immunodominant T cell epitopes, aligning with the earlier observed high IL-17a induction in the pre-challenge spleen (**Fig. 4B**). These results demonstrate that BECC adjuvants significantly improve immune recognition of linear HA epitopes, particularly those localized to the protein’s head domain. Notably, BECC470s and BECC470b are highly effective in stimulating a broad and robust antibody response against multiple epitopes of the HA protein.

**Fig. 6.**
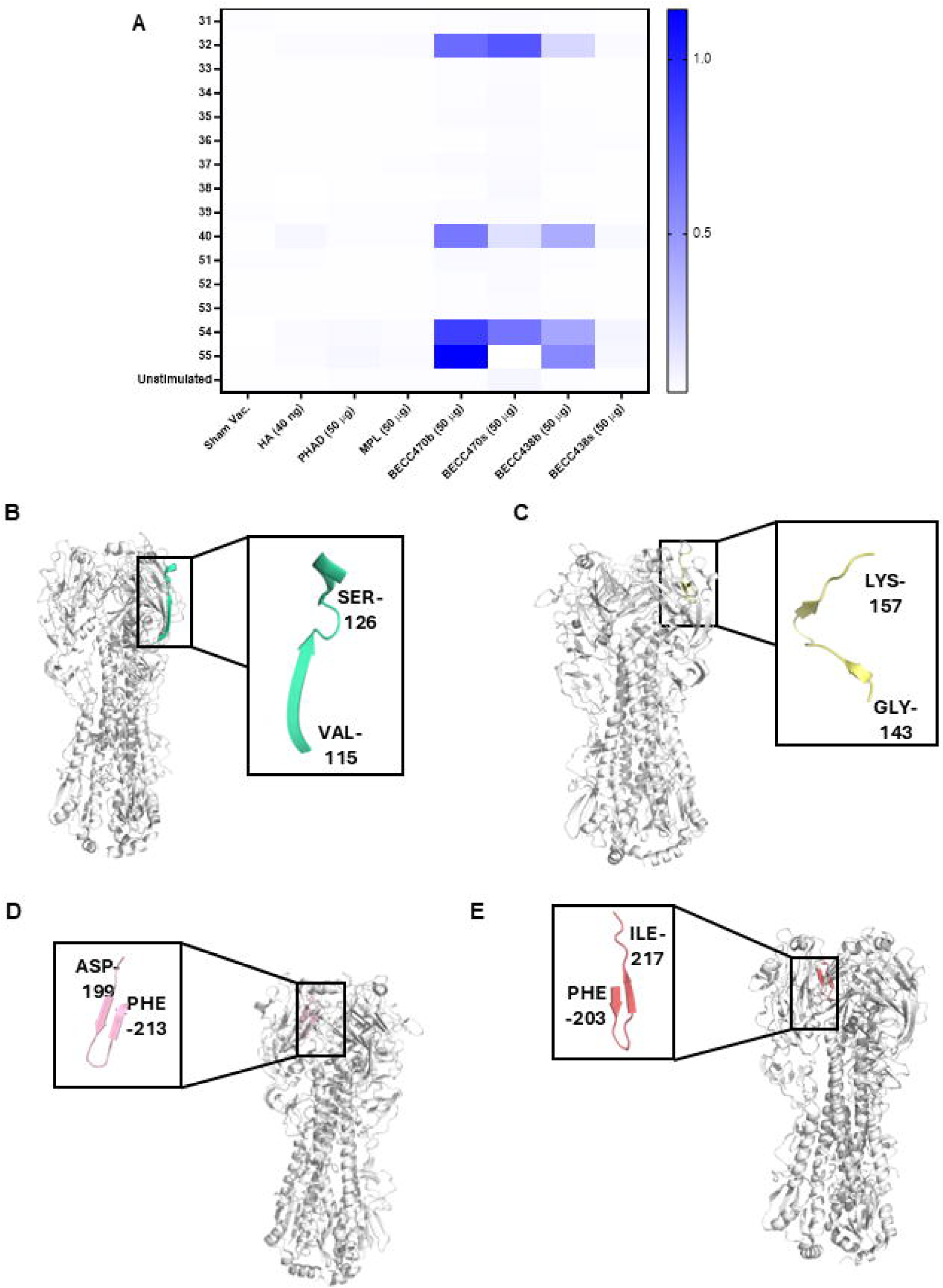
Identification of linear T cell epitopes. Six-week-old BALB/c mice (n=5/group) were immunized intramuscularly on days 0 and 14 with 40 ng of rHA with indicated adjuvants. Sham groups were immunized with PBS. On day 35 post-immunization, mice were sacrificed and single cell suspensions from spleens were prepared. Cells were plated in 96 well plates and stimulated with individual peptides. IL-17a secretion in supernatants was determined, and the data was plotted as heat map to visualize immunodominant T cell epitopes (**A**). **B-E** shows the individual linear T cell epitopes while **D** and **E** have partially overlapped sequences.

## Induction of immunological memory by BECC’s

To assess the potential for long-term immunity, we investigated the ability of BECC-adjuvanted groups to generate memory B cells, a key component of immunological memory. Mice were immunized as previously described, and spleens were harvested on day 35 post-immunization. Single-cell suspensions were cultured for 5 days with mitogen to facilitate the conversion of memory B cells into effector B cells, with cells cultured without mitogens serving as controls. ELISpot assays were then performed to detect HA-specific IgG and IgA-secreting cells. The results showed that BECC470b, followed closely by BECC470s, induced the highest number of IgG-secreting memory B cells, significantly surpassing the levels observed in groups adjuvanted with MPL (**Fig. 7**). However, only BECC470b demonstrated a significantly higher level of IgG-secreting memory B cells as compared to the MPL-adjuvanted group. No IgA-secreting memory B cells were detected in the spleens of any group. These findings demonstrate that BECC470b and BECC470s effectively induce memory B cell formation, may effectively promote long-lasting immunity through the generation of memory B cells.

**Fig. 7.**
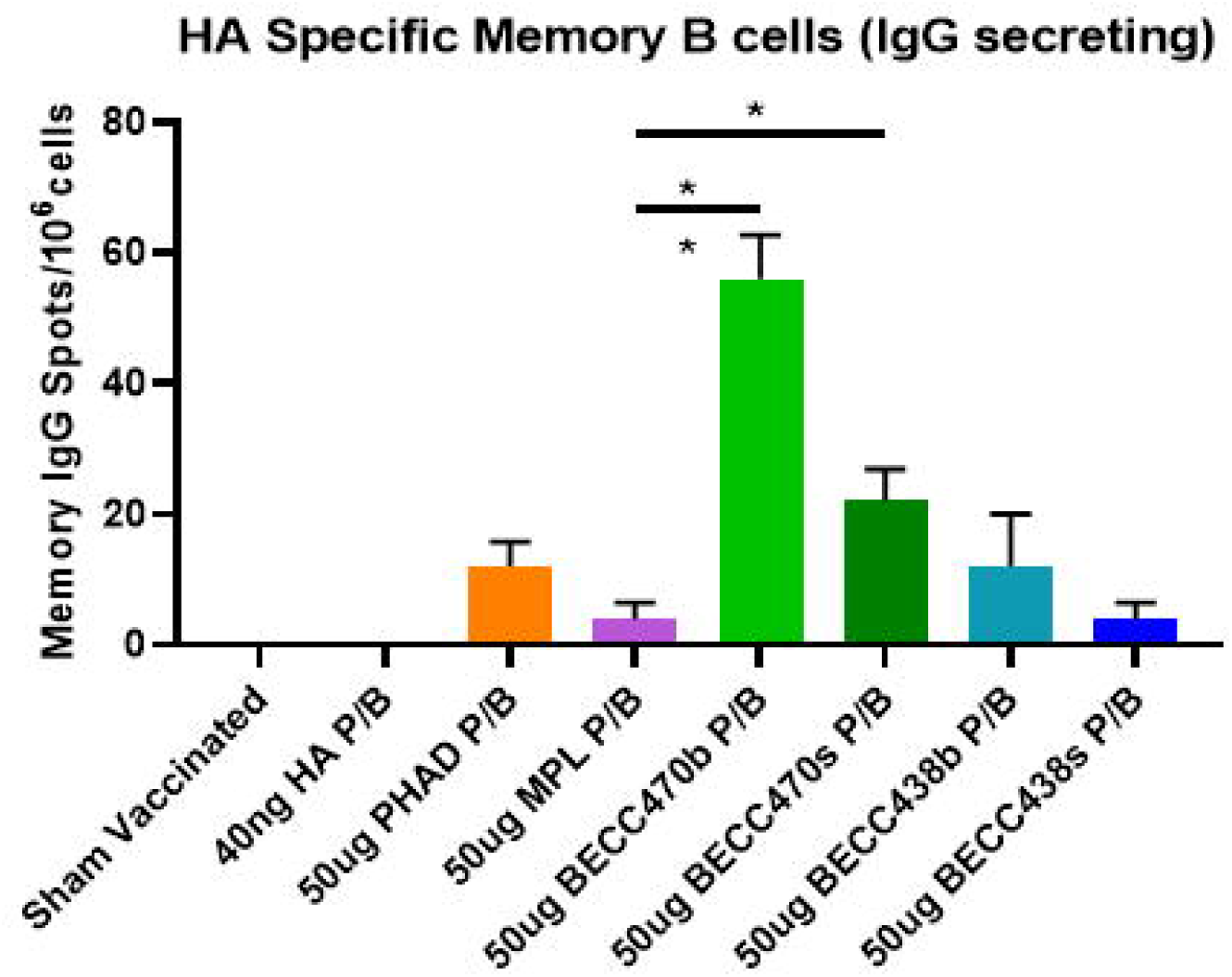
Induction of memory B cells responses by BECC adjuvanted rHA. Six-week-old BALB/c mice (n=5/group) were immunized intramuscularly on days 0 and 14 with 40 ng of rHA with indicated adjuvants. Sham groups were immunized with PBS. On day 35 post-immunization, mice were sacrificed and single cell suspensions from spleens were prepared. Memory B cell was detected by ELISpot as described in materials and methods. Data were analyzed by Dunnett’s multiple comparison test by comparing to MPL adjuvanted group. *p<0.05, **p<0.01, ***p<0.001.

## Durability of immune response generated by BECC’s

Sustaining long-term immunity without booster doses represents a critical hallmark of effective vaccine formulations. To evaluate the capacity of BECC molecules to induce a durable immune response, mice were vaccinated using the previously described protocol above. Seventeen months post-vaccination, intranasal challenge with 1000 PFU of influenza A/Netherlands/02/09 (H1N1) revealed that BECC470s-and BECC438b-immunized mice maintained superior protection during 7-day monitoring (**Fig. 8**), while all other groups showed ≥15% weight reduction. These findings demonstrate that BECC470s and BECC438b can induce a durable immune response that provides long-term protection against influenza infection, even after a prolonged period post-vaccination.

**Fig. 8.**
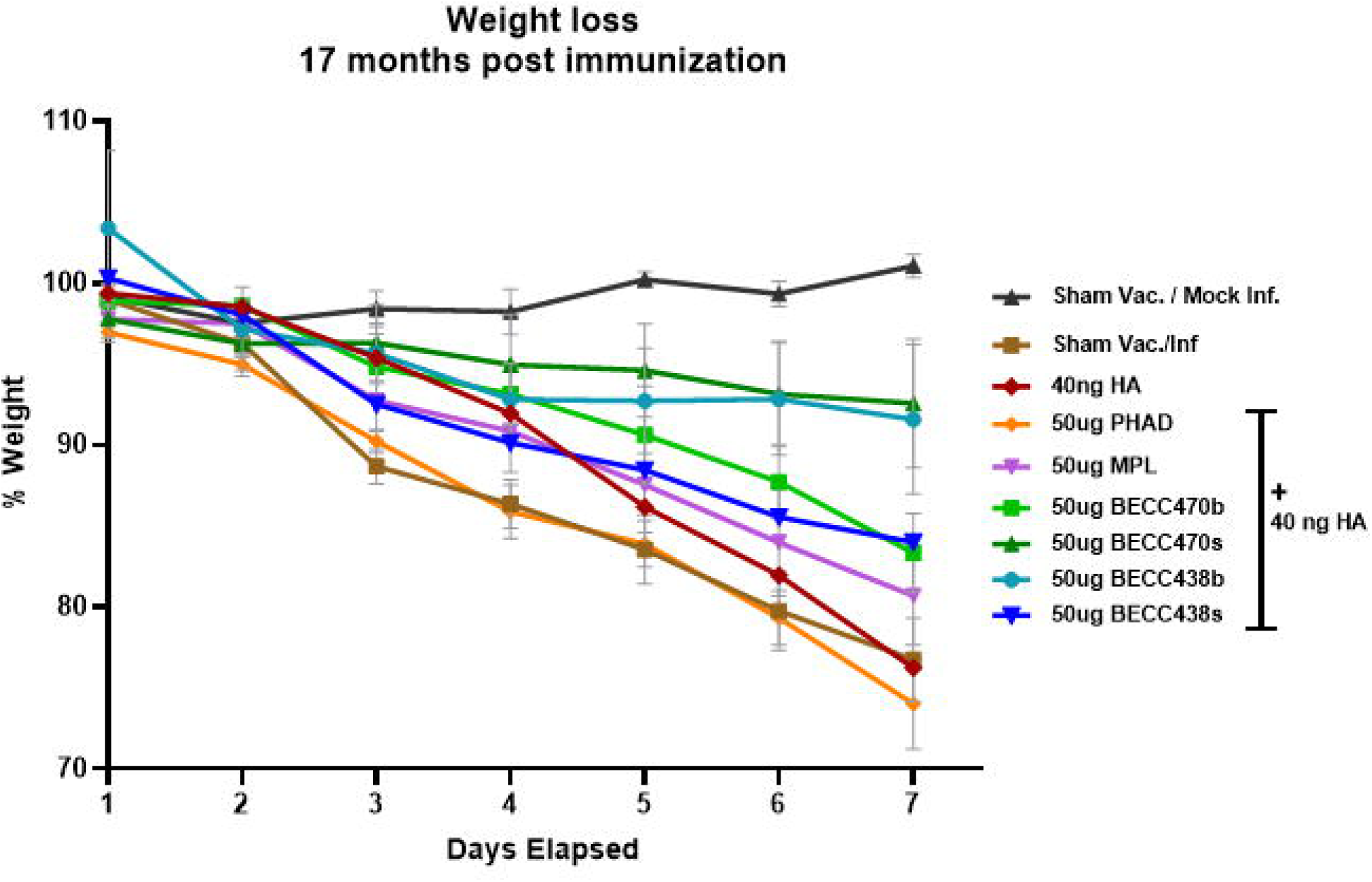
Durability of immune response generated by BECCs. Six-week-old BALB/c mice were immunized intramuscularly on days 0 and 14 with 40 ng of rHA with indicated adjuvants. Sham groups were immunized with PBS. Mice were intranasally challenged with 3200 PFU of NL09 (H1N1) on day 35 post-immunization. Body weights were monitored for the infected mice.

## Discussion

The 2023-24 influenza season has provided valuable insights into vaccine effectiveness (VE) across different age groups and influenza strains. For IAV, children and adolescents under 18 years showed higher protection, with VE ranging from 59% to 67% against outpatient visits. Adults over 18 years experienced more modest protection, with VE between 33% and 49%. Influenza B virus vaccines demonstrated higher effectiveness across all age groups, with VE estimates of 64% to 89% for pediatric patients and 60% to 78% for adults ^16^. These findings highlight the differential impact of current vaccine formulations on various demographic groups and influenza strains.

The availability of eight quadrivalent vaccines for the current season, including two specifically designed for individuals over 65 years, reflects efforts to address the diverse needs of the population. One of these specialized vaccines contains a high dose of hemagglutinin (60 micrograms of each HA), while the other is adjuvanted with MF59, a squalene-based oil-in-water emulsion. Despite these advancements and the range of available options, the overall VE can be described as modest, underscoring the ongoing challenge of developing more effective influenza vaccines. This situation emphasizes the need for continued research and development to enhance vaccine efficacy and provide better protection against influenza across all age groups^17,18^.

Adjuvants are incorporated into vaccine formulations to strengthen immune responses and counteract suboptimal vaccine effectiveness. Aluminum salts (potassium aluminum sulfate and aluminum hydroxide) have been traditionally used as adjuvants in various FDA-approved vaccines^19,20^. More recently, oil-in-water emulsions have been employed as adjuvants in the H5N1 (bird flu) vaccine (Aflunov) and the quadrivalent influenza vaccine formulated for older adults (Fluad) ^21^. The shingles vaccine (Shingrix) is formulated with AS01B, while the RSV vaccine (AREXVY) uses AS01E; both formulations include MPL and QS-21, a saponin derived from the bark of the Chilian soap bark tree (*Quillaja saponaria*). Although MPL is well-characterized, its biological origin results in multiple congeners of lipid A, which can introduce variability in immune responses. To mitigate this variability, a synthetic version of MPL called phosphorylated hexa-acyl disaccharide (PHAD) was developed and tested for efficacy in clinical trials ^22–24^.

In response to the need for improved adjuvants, Bacterial Enzymatic Combinatorial Chemistry (BECC) has been employed to create lead biologically-derived TLR4 agonists, BECC438b and BECC470b^25^. These adjuvants have shown efficacy against a diverse range of bacterial and viral pathogens, including *Yersinia, Pseudomonas, Salmonella, Shigella,* HPV, influenza A, and SARS-CoV-2 ^8,13–15,26–28^. In addition to their biological forms, synthetic versions (BECC438s and BECC470s) have also been developed. Structurally, both BECC438s and BECC470s are symmetrical (3 + 3)-hexa-acylated molecules (**Fig. 1**) containing two secondary C16 acyl chains added at the 2 and 2′ positions. The acyl chain at the 2′ position has one unsaturation between the ninth and tenth carbon. In contrast, BECC438s is bis-phosphorylated while BECC470s is mono-phosphorylated and lacks the 4’ phosphate moiety. These structural distinctions may significantly influence the immunological properties and overall effectiveness of these adjuvants.

To investigate the impact of unique BECC structures on the efficacy and immunogenicity of the antigen, both biological and synthetic versions of BECC molecules were tested in a mouse IAV challenge model using rHA as the antigen. Previously, BECC470b has demonstrated antigen-sparing properties, and in this study, we utilized 40 ng of trimeric rHA derived from H1N1 (Cal09)^14^. The serum IgG1 and IgG2a responses were negligible with the protein alone, showed modest improvement with comparator adjuvants (PHAD and MPL), and increased dramatically with the BECC adjuvants. Notably, IgG1 levels, which are known to facilitate the neutralization of viral particles, were significantly higher across all BECC-immunized groups, likely contributing to the lower viral loads observed in their lungs compared to those receiving MPL and PHAD adjuvants. Additionally, groups adjuvanted with BECC470 exhibited enhanced viral clearance and elevated levels of IgG2a induction as compared with control adjuvants. This finding is supported by research indicating that influenza vaccines that induce higher levels of IgG2a tend to be more efficacious, as mice that survive viral infections typically demonstrate increased levels of this isotype ^9^. Although IgG2a is not directly responsible for neutralizing the virus, it plays a crucial role in promoting viral clearance ^9,29,30^. This underscores the importance of IgG2a in the overall immune response to influenza, highlighting its potential contribution to the effectiveness of vaccines formulated with BECC470 in mice.

We next compared the cell-mediated immune response generated by the adjuvants and found that splenocytes isolated from mice adjuvanted with BECC exhibited significantly higher levels of IL-17A, IL-12p70, IFN-γ, and IL-6 secretion compared to those from control adjuvants. Similar but elevated levels of these cytokines were observed in cells isolated from the lungs of mice challenged with NL09 (H1N1), indicating that systemic immune cells migrate to the lungs, the primary site of infection, upon challenge. The pro-inflammatory cytokines induced by BECC have been reported to have both beneficial and detrimental effects during influenza infections ^31^; for instance, while neutralization of IL-17A can reduce lung injury during infection, IL-17A-deficient mice struggle to clear viral infections due to impaired antibody production from B-1a cells in the pulmonary tract ^32^. Interestingly, IL-17A has also been found to prevent secondary bacterial pneumonia induced by *Streptococcus* following influenza infections, suggesting that a desirable side effect of BECC-induced IL-17A may be its ability to prevent such secondary infections ^33^.

After investigating the humoral and cell-mediated responses to the whole rHA protein, we mapped linear epitopes to gain a deeper understanding of the immunomodulation provided by the adjuvants. Linear B cell epitope analysis indicated that BECC adjuvants, particularly BECC470 (b and s), significantly enhanced immune recognition of immunogenic epitopes, which likely accounts for the higher antibody titers observed against the full-length HA. Additionally, linear T cell epitope analysis may clarify why BECC adjuvants, especially BECC470b and BECC470s, exhibit superior viral clearance capabilities ^34^; similar to their effects on B cells, these adjuvants increased immune recognition of linear T cell epitopes. Notably, both BECC470b and BECC438b were able to elicit immune recognition against a specific peptide (#55), which was absent in all other adjuvants. Physiologically, antibodies targeting the head region of HA have been shown to effectively inhibit receptor engagement and prevent infection^35,36^. These observations suggest that BECC adjuvants enhance immune recognition of linear HA epitopes which are located on the head region of the protein. These data indicates that BECC adjuvants, particularly BECC470s and BECC470b, are highly effective in stimulating a robust antibody response against multiple epitopes of the rHA protein. This enhanced recognition of linear epitopes may contribute to the superior protection observed in BECC-adjuvanted groups, potentially by targeting a wider range of viral epitopes and improving the overall quality of the immune response. Furthermore, both versions of BECC470 can induce memory B cells, which can quickly differentiate into antibody-secreting B cells upon encountering an infection ^37^.

In summary, the development of BECC adjuvants addresses the pressing need for more effective vaccine formulations that can enhance immune responses. These adjuvants have demonstrated the ability to stimulate a balanced Th1/Th2 immune response, which is essential for providing comprehensive protection against infections^38^. By utilizing the BECC platform, novel adjuvants can be produced in a reproducible and cost-effective manner, making them suitable for large-scale vaccine production. The promising results from studies involving BECC438 and BECC470 underscore their potential as superior adjuvants in combination with vaccines targeting various pathogens, paving the way for improved vaccination strategies in the future. Current research is also exploring various formulations, such as AS0-biosimilars, to further augment immune responses, along with investigating the role of these formulations in adjuvant sparing.

## Materials and Methods

### Study Design

This study aimed to compare the adjuvanticity of biological and synthetic versions of BECC438, BECC470, MPL, and PHAD using rHA as an antigen in a mouse model of influenza infection. Mice were randomly assigned treatment groups. To assess protection, body weight loss was assessed in vaccinated mice post-challenge with influenza virus. To assess humoral immune response, immune sera was evaluated using ELISA. Pre and post challenge cytokine response was evaluated in single cell suspensions of spleens and lungs using multiplex assay. Immunogenic linear epitopes were identified using peptide arrays and memory B cells were assayed in spleens of immunized mice by ELISpot. Sample size was chosen based on prior experience and is mentioned in respective figure legends. All data points, including outliers, if any, were considered. Experiments were repeated at least twice, and representative data is shown.

### Protein production

Influenza rHA from H1 A/California/07/2009 was produced as previously described ^39^. Briefly, to facilitate secretion and purification from the supernatant of Expi293F cells, the transmembrane domain sequence was replaced with a heterologous carboxy (C)-terminal trimerization domain (i.e., fibritin foldon), or with the GCN4 isoleucine zipper trimerization domain from Saccharomyces cerevisiae, both with a C-terminal 6X-histidine (6XHIS) tag. Recombinant HAs were verified by performing ELISAs using a panel of previously described mAbs that recognize conformational epitopes on the rHA stalk, as described ^39^.

### Mice and Immunizations

Six-to eight-week-old, BALB/c mice (Jackson Laboratories, ME, USA) were immunized intramuscularly (IM) in the hind, caudal thigh at day 0 (prime) and day 14 (boost). Fifty µL of vaccine solution were prepared by admixing 0.04 µg of rHA with a fibritin trimerization domain (Influenza/A/California/04/2009) and 50 µg of either BECC470b (or synthetic “s”), BECC438b (or synthetic “s”), MPL (Invivogen, CA, USA), and PHAD (Croda, AL, USA) was used. The vaccines were incubated with rocking for 1 hour to allow interaction between the antigen and the adjuvants. Blood was collected by saphenous vein on days 14, 28, and at time of harvest. Animals were euthanized in compliance with AVMA guideline. Briefly, using a non-pre-charged chamber, carbon dioxide gas was dispensed from a commercial cylinder with fixed pressure regulator and inline restrictor controlling the gas flow within 30-70 % of the chamber volume per minute. After respirator arrest, gas flow as maintained for at least 60 seconds, followed by bilateral thoracotomy or cervical dislocation, to ensure euthanasia. All animal procedures were reviewed and approved by University of Maryland, Baltimore Institutional Animal Care and Use Committee.

### Influenza Infection

35 days post-immunizations, mice were sedated by intraperitoneal administration of ketamine/xylene and intranasally inoculated groups of mice with 3200 plaque forming units of homologous A/Netherlands/602/2009 (H1N1) (NL09). Daily weights were recorded post challenge. Mice that fell below 70 percent of initial body weight were humanely euthanized in accordance with IACUC protocols.

### ELISA

Antibodies specific for rHA were determined by ELISA as described previously with minor modifications ^8^. Briefly, 96-well plates (Nunc, NY, USA) were coated with rHA containing GCN4 trimerization domain (2 µg/mL in 0.1 M sodium bicarbonate buffer solution) overnight at 4°C. Plates were washed with wash buffer (PBS-Tween 20 (0.05%)) and blocked for two hours at room temperature with blocking buffer (0.5% nonfat dry milk, 3% goat serum in wash buffer). Blocking buffer was discarded, serum sample dilutions were added, and plates incubated for 2 hours at room temperature. After washing, HRP conjugated secondary antibody (1:3000; Thermo Scientific, MA, USA) against IgG, IgG1 and IgG2a were added to separate plates and incubated for 1 hour at room temperature. Post washing, 100 μl of 3,3′,5,5′-Tetramethylbenzidine (BD Biosciences, CA, USA) substrate was added, and reaction was stopped with an equal volume of stop solution (KPL, MD, USA) after 15 minutes. Titers were calculated using GraphPad prism (GraphPad Software, MA, USA).

### Isolation of murine splenocytes and lungs

Spleens and lungs were harvested from euthanized mice and placed in 1 ml of serum-free RMPI immediately on ice. Spleens and lungs were dissociated using the gentleMACS Octo Dissociator in gentleMACS (Miltenyi Biotec, CA, USA) according to manufacturer’s instructions. Briefly, the organs were transferred to a c-tube containing an enzyme mixture and loaded onto the instrument for dissociation. After completion of the cycle, dissociated organ mixture was filtered using MACS SmartStrainers (80 µM) [Miltenyi Biotec, CA, USA], pelleted at 800 x g for 5 minutes, and supernatants discarded. Cell pellets were resuspended in red blood cell lysis buffer and incubated for seven minutes, and the reaction was stopped by adding 8 ml of complete RPMI (Thermo Scientific, MA, USA). The cells were centrifuged (800 x g, 5 minutes), resuspended in complete RPMI, counted, and stored on ice until further processing.

### Cytokine determinations

Cells isolated from lung and spleen were counted and incubated with 10 µg/ml rHA with GCN4 trimerization or PBS in 96 well plates (10^6^ cell/well) for 48 h at 37°C. Supernatants were collected and analyzed for cytokines: IFN-γ, IL-10, IL-12p70, IL-17A, IL-1β, IL-2, IL-4, IL-5, IL-6, and TNF-α with concentrations being determined using an MSD plate reader with associated analytical software (Meso Scale Discovery, MD, USA). While multiple cytokines were measured, only the ones that were differentially modulated upon stimulation are reported here.

### B cell linear epitope mapping

Peptide array for HA (H1 A/California/07/2009) was obtained from BEI Resources, NIAID, NIH (NR-19244). The 139 peptide arrays (15-mers with 11mer overlaps) were dissolved at a concentration of 1 mg/ml using manufacturer suggested solvents. 10 sequential peptides were mixed to create 14 peptide pools. Peptides were covalently linked by the free carboxylic acid group to amino group of the 96 well plate by coupling agent 1-Ethyl-3-(3-dimethylaminopropyl)carbodiimide (EDC, Sigma, MO, USA). Briefly, pools were diluted with MES buffer (pH 5.5) (Sigma, MO, USA) at a final concentration of 0.1 mg/ml. The solution was added to 96 well plates followed by 10 μl of EDC (10mg/ml in water). Plates were incubated for 1 hour at room temperature and washed with distilled water. Reactivity against immune serum was determined as described in ELISA. To evaluate the reactivity against individual peptides in the pool, each peptide was diluted in MES buffer and coupled onto plates as described earlier.

### T cell epitope mapping

Splenocytes from immunized mice were isolated as described earlier and simulated with peptide the peptide pools from B cell linear epitope mapping experiment. 48 hours post immunization, supernatants were collected and analyzed for IL-17a secretion using direct ELISA (DuoSet, R&D Systems, MN, USA) following manufacturer’s protocol. Individual peptides from pools that were positive for IL-17a secretion were used to stimulate splenocytes isolated from immunized mice. IL-17a secretion in supernatants were analyzed to identify T cell epitopes.

### Memory B cell

To detect HA-specific memory B cells, splenocytes isolated from immunized mice were cultured in complete RPMI containing mitogen B-Poly-S (C.T.L., OH, USA) in 5% CO_2_ at 37°C for 4 days. Untreated cells were cultured to be used as control. After 4 days, cells were washed with PBS, resuspended in complete RPMI, counted, equalized and ELISpot was performed according to manufacturer’s protocol (CTL, OH, USA) to determine HA-specific memory B cells. Briefly, rHA with GCN4 trimerization domain (5 μg/ml) in PBS was added to pre-wetted membrane bottom 96 well plates and incubated at 4°C overnight. Plates were washed with PBS and blocked with complete RPMI at 37°C for 1 hour. Blocking media was discarded, cells added, and plates were incubated in 5% CO_2_ at 37°C for 18 hours. Cells were discarded, plates washed with PBS and detection solution was added and incubated at room temperature for 2 hours. Following washing, tertiary solution was added and plates were incubated at room temperature for 1 hour. Spots were developed using provided substrates and the reaction was stopped by washing plates in tap water. The plates were dried overnight and scanned using CTL Immunospot reader. Data was represented as spots/10^6^ cells. Spot obtained in non-mitogen stimulated wells were subtracted from their respective mitogen stimulated counterparts.

### Statistical analysis

GraphPad Prism 10.4.1 (MA, USA) was used to perform all statistical comparisons. Differences among treatment groups were analyzed as mentioned in figure legends. A p-value of less than 0.05 was considered significant for all comparisons. * P< 0.05; ** P<0.01; *** P< 0.001.

### List of Supplementary Materials

This pdf file includes:

Figs. S1 to S3 Table S1

## Acknowledgements

The following reagent was obtained through BEI Resources, NIAID, NIH: Peptide Array, Influenza Virus A/California/07/2009 (H1N1)pdm09 Hemagglutinin Protein, NR-19244. The authors are grateful to Charles Richardson, and Hans Lien for generous critical review of this work.

## Author Contributions

Conceptualization: SD, FMG, RKE

Methodology: SD, BMT, MBF, RKE

Investigation: SD, BMT, DR, SS, LB, DJV

MS analysis: HY

Reagents (rHA): HN, LK, FK, LC

Reagents (H1N1 virus): WS

Funding acquisition: SD, FMG, RKE

Project administration: FMG

Writing – original draft: SD

Writing – review & editing: SD, BMT, FMG, MBF, RKE

All authors read and approved the manuscript.

## Data and materials availability

All data are available in the main text or the supplementary materials; further inquiries can be directed to the corresponding author.

## References

1 Shichinohe, S. & Watanabe, T. Advances in Adjuvanted Influenza Vaccines. Vaccines (Basel) 11 (2023). 10.3390/vaccines11081391

2 Harrington, W. N., Kackos, C. M. & Webby, R. J. The evolution and future of influenza pandemic preparedness. Exp Mol Med 53, 737–749 (2021). 10.1038/s12276-021-00603-0

3 Krammer, F. The human antibody response to influenza A virus infection and vaccination. Nat Rev Immunol 19, 383–397 (2019). 10.1038/s41577-019-0143-6

4 Coleman, B. L., Sanderson, R., Haag, M. D. M. & McGovern, I. Effectiveness of the MF59-adjuvanted trivalent or quadrivalent seasonal influenza vaccine among adults 65 years of age or older, a systematic review and meta-analysis. Influenza Other Respir Viruses 15, 813–823 (2021). 10.1111/irv.12871

5 Pulendran, B. P. S. A. & O’Hagan, D. T. Emerging concepts in the science of vaccine adjuvants. Nat Rev Drug Discov 20, 454–475 (2021). 10.1038/s41573-021-00163-y

6 Facciola, A., Visalli, G., Lagana, A. & Di Pietro, A. An Overview of Vaccine Adjuvants: Current Evidence and Future Perspectives. Vaccines (Basel) 10 (2022). 10.3390/vaccines10050819

7 Tregoning, J. S., Russell, R. F. & Kinnear, E. Adjuvanted influenza vaccines. Hum Vaccin Immunother 14, 550–564 (2018). 10.1080/21645515.2017.1415684

8 Haupt, R. E. et al. Novel TLR4 adjuvant elicits protection against homologous and heterologous Influenza A infection. Vaccine 39, 5205–5213 (2021). 10.1016/j.vaccine.2021.06.085

9 Huber, V. C. et al. Distinct contributions of vaccine-induced immunoglobulin G1 (IgG1) and IgG2a antibodies to protective immunity against influenza. Clin Vaccine Immunol 13, 981–990 (2006). 10.1128/CVI.00156-06

10 Baldwin, S. L. et al. The importance of adjuvant formulation in the development of a tuberculosis vaccine. J Immunol 188, 2189–2197 (2012). 10.4049/jimmunol.1102696

11 Bertholet, S. et al. Optimized subunit vaccine protects against experimental leishmaniasis. Vaccine 27, 7036–7045 (2009). 10.1016/j.vaccine.2009.09.066

12 Haupt, R. et al. Enhancing the protection of influenza virus vaccines with BECC TLR4 adjuvant in aged mice. Sci Rep 13, 715 (2023). 10.1038/s41598-023-27965-x

13 Howlader, D. R. et al. Effect of Two Unique Nanoparticle Formulations on the Efficacy of a Broadly Protective Vaccine Against Pseudomonas Aeruginosa. Front Pharmacol 12, 706157 (2021). 10.3389/fphar.2021.706157

14 Lu, T. et al. Impact of the TLR4 agonist BECC438 on a novel vaccine formulation against Shigella spp. Front Immunol 14, 1194912 (2023). 10.3389/fimmu.2023.1194912

15 Das, S. et al. Immunogenicity and protective efficacy of nanoparticle formulations of L-SseB against Salmonella infection. Front Immunol 14, 1208848 (2023). 10.3389/fimmu.2023.1208848

16 Ruth Link-Gelles, P. S. C., DHSc2; Alexander Webber, MPH1; Toan C. Ong, PhD3; Elizabeth A.K. Rowley, DrPH2; Malini B. DeSilva, MD4; Kristin Dascomb, MD, PhD5; Stephanie A. Irving, MHS6; Nicola P. Klein, MD, PhD7; Shaun J. Grannis, MD8,9; Michelle A. Barron3; Sarah E. Reese, PhD2; Charlene McEvoy, MD4; Tamara Sheffield, MD5; Allison L. Naleway, PhD6; Ousseny Zerbo, PhD7; Colin Rogerson, MD9,10; Wesley H. Self, MD11; Yuwei Zhu, MD11; Adam S. Lauring, MD, PhD12; Emily T. Martin, PhD12; Ithan D. Peltan, MD13,14; Adit A. Ginde, MD15; Nicholas M. Mohr, MD16; Kevin W. Gibbs, MD17; David N. Hager, MD, PhD18; Matthew E. Prekker, MD19; Amira Mohamed, MD20; Nicholas Johnson, MD21; Jay S. Steingrub, MD22; Akram Khan, MBBS23; Jamie R. Felzer, MD24; Abhijit Duggal, MD25; Jennifer G. Wilson, MD26; Nida Qadir, MD27; Christopher Mallow, MD28; Jennie H. Kwon, DO29; Cristie Columbus, MD30,31; Ivana A. Vaughn, PhD32; Basmah Safdar, MD33; Jarrod M. Mosier, MD34; Estelle S. Harris, MD14; James D. Chappell, MD, PhD11; Natasha Halasa, MD11; Cassandra Johnson, MS11; Karthik Natarajan, PhD35,36; Nathaniel M. Lewis, PhD37; Sascha Ellington, PhD37; Emily L. Reeves, MPH37; Jennifer DeCuir, MD, PhD37; Meredith McMorrow, MD1; Clinton R. Paden, PhD1; Amanda B. Payne, PhD1; Fatimah S. Dawood, MD1; Diya Surie, MD1. Interim Estimates of 2024–2025 COVID-19 Vaccine Effectiveness Among Adults Aged ≥18 Years — VISION and IVY Networks, September 2024–January 2025. 73-82 (2024).

17 Trombetta, C. M., Kistner, O., Montomoli, E., Viviani, S. & Marchi, S. Influenza Viruses and Vaccines: The Role of Vaccine Effectiveness Studies for Evaluation of the Benefits of Influenza Vaccines. Vaccines (Basel) 10 (2022). 10.3390/vaccines10050714

18 Taaffe, J. et al. Advancing influenza vaccines: A review of next-generation candidates and their potential for global health impact. Vaccine 42, 126408 (2024). 10.1016/j.vaccine.2024.126408

19 Marrack, P., McKee, A. S. & Munks, M. W. Towards an understanding of the adjuvant action of aluminium. Nat Rev Immunol 9, 287–293 (2009). 10.1038/nri2510

20 HogenEsch, H., O’Hagan, D. T. & Fox, C. B. Optimizing the utilization of aluminum adjuvants in vaccines: you might just get what you want. NPJ Vaccines 3, 51 (2018). 10.1038/s41541-018-0089-x

21 Clegg, C. H., Rininger, J. A. & Baldwin, S. L. Clinical vaccine development for H5N1 influenza. Expert Rev Vaccines 12, 767–777 (2013). 10.1586/14760584.2013.811178

22 Falloon, J. et al. A phase 1a, first-in-human, randomized study of a respiratory syncytial virus F protein vaccine with and without a toll-like receptor-4 agonist and stable emulsion adjuvant. Vaccine 34, 2847–2854 (2016). 10.1016/j.vaccine.2016.04.002

23 Heeke, D. S. et al. Identification of GLA/SE as an effective adjuvant for the induction of robust humoral and cell-mediated immune responses to EBV-gp350 in mice and rabbits. Vaccine 34, 2562–2569 (2016). 10.1016/j.vaccine.2016.04.012

24 Pantel, A. et al. A new synthetic TLR4 agonist, GLA, allows dendritic cells targeted with antigen to elicit Th1 T-cell immunity in vivo. Eur J Immunol 42, 101–109 (2012). 10.1002/eji.201141855

25 Gregg, K. A. et al. Rationally Designed TLR4 Ligands for Vaccine Adjuvant Discovery. mBio 8 (2017). 10.1128/mBio.00492-17

26 Gregg, K. A. et al. A lipid A-based TLR4 mimetic effectively adjuvants a Yersinia pestis rF-V1 subunit vaccine in a murine challenge model. Vaccine 36, 4023–4031 (2018). 10.1016/j.vaccine.2018.05.101

27 Matthews, R. L. et al. Immune profile diversity is achieved with synthetic TLR4 agonists combined with the RG1-VLP vaccine in mice. Vaccine 44, 126577 (2025). 10.1016/j.vaccine.2024.126577

28 Lee, K. S. et al. Intranasal VLP-RBD vaccine adjuvanted with BECC470 confers immunity against Delta SARS-CoV-2 challenge in K18-hACE2-mice. Vaccine 41, 5003–5017 (2023). 10.1016/j.vaccine.2023.06.080

29 Rubtsova, K., Rubtsov, A. V., van Dyk, L. F., Kappler, J. W. & Marrack, P. T-box transcription factor T-bet, a key player in a unique type of B-cell activation essential for effective viral clearance. Proc Natl Acad Sci U S A 110, E3216–3224 (2013). 10.1073/pnas.1312348110

30 Markine-Goriaynoff, D. & Coutelier, J. P. Increased efficacy of the immunoglobulin G2a subclass in antibody-mediated protection against lactate dehydrogenase-elevating virus-induced polioencephalomyelitis revealed with switch mutants. J Virol 76, 432–435 (2002). 10.1128/jvi.76.1.432-435.2002

31 Ma, W. T., Yao, X. T., Peng, Q. & Chen, D. K. The protective and pathogenic roles of IL-17 in viral infections: friend or foe? Open Biol 9, 190109 (2019). 10.1098/rsob.190109

32 Wang, X. et al. IL-17A Promotes Pulmonary B-1a Cell Differentiation via Induction of Blimp-1 Expression during Influenza Virus Infection. PLoS Pathog 12, e1005367 (2016). 10.1371/journal.ppat.1005367

33 Loughran, S. T. et al. Influenza infection directly alters innate IL-23 and IL-12p70 and subsequent IL-17A and IFN-gamma responses to pneumococcus in vitro in human monocytes. PLoS One 13, e0203521 (2018). 10.1371/journal.pone.0203521

34 Schroeder, S. M., Nelde, A. & Walz, J. S. Viral T-cell epitopes - Identification, characterization and clinical application. Semin Immunol 66, 101725 (2023). 10.1016/j.smim.2023.101725

35 Leon, P. E. et al. Optimal activation of Fc-mediated effector functions by influenza virus hemagglutinin antibodies requires two points of contact. Proc Natl Acad Sci U S A 113, E5944–E5951 (2016). 10.1073/pnas.1613225113

36 Zost, S. J. et al. Identification of Antibodies Targeting the H3N2 Hemagglutinin Receptor Binding Site following Vaccination of Humans. Cell Rep 29, 4460–4470 e4468 (2019). 10.1016/j.celrep.2019.11.084

37 Wong, R. & Bhattacharya, D. Basics of memory B-cell responses: lessons from and for the real world. Immunology 156, 120–129 (2019). 10.1111/imm.13019

38 Aleebrahim-Dehkordi, E. et al. T helper type (Th1/Th2) responses to SARS-CoV-2 and influenza A (H1N1) virus: From cytokines produced to immune responses. Transpl Immunol 70, 101495 (2022). 10.1016/j.trim.2021.101495

39 Bliss, C. M. et al. A single-shot adenoviral vaccine provides hemagglutinin stalk-mediated protection against heterosubtypic influenza challenge in mice. Mol Ther 30, 2024–2047 (2022). 10.1016/j.ymthe.2022.01.011

