## Supplemental for "Enhancing protective efficacy and immunogenicity of hemagglutinin-based influenza vaccine utilizing adjuvants developed by BECC"

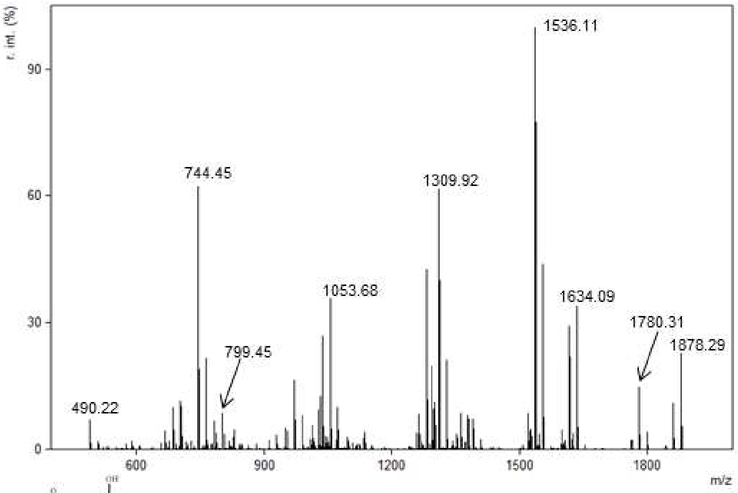

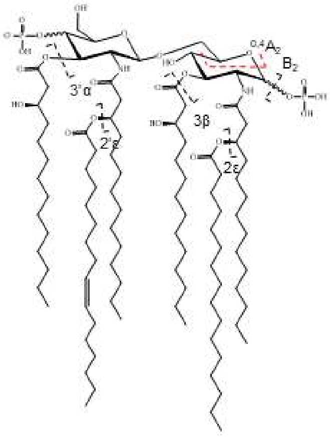

| Naming | *m/z* (theory) | *m/z* (expt) | Error (Da) |
|---|---|---|---|
| $B_2$ | 1780.31 | 1780.29 | 0.02 |
| 3'α | 1634.09 | 1634.07 | 0.02 |
| $B_2$ + 3'α | 1536.11 | 1536.09 | 0.02 |
| $B_2$ + 3'α + 3β | 1309.92 | 1309.90 | 0.02 |
| $B_2$ + 3'α + 3β + 2ε | 1053.68 | 1053.67 | 0.01 |
| $B_2$ + 3'α + 3β + 2ε + 2'ε | 799.45 | 799.44 | 0.01 |
| 3'α + $^{0,4}A_2$ | 744.45 | 744.44 | 0.01 |
| 3'α + 2'ε + $^{0,4}A_2$ | 490.22 | 490.22 | 0.00 |

**BECC438s**
Chemical Formula: $C_{100}H_{187}N_2O_{25}P_2^-$
Exact Mass: 1878.29

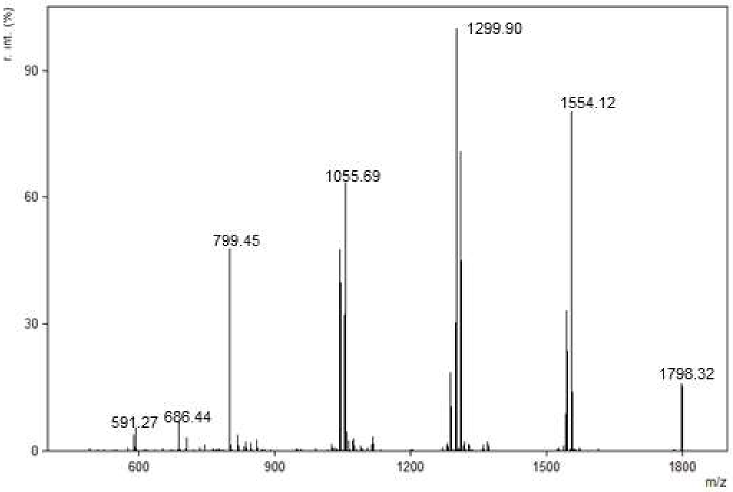

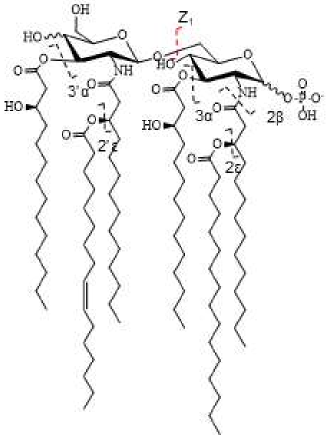

| Naming | *m/z* (theory) | *m/z* (expt) | Error (Da) |
|---|---|---|---|
| 3'α | 1554.12 | 1554.10 | 0.02 |
| 3'α + 2'ε | 1299.90 | 1299.88 | 0.02 |
| 3'α + 3α + 2ε | 1055.69 | 1055.68 | 0.01 |
| 3'α + 3β + 2ε + 2'ε | 799.45 | 799.44 | 0.01 |
| 3α + $Z_1$ | 686.44 | 686.44 | 0.00 |
| 3'α + 3β + 2ε + 2'ε + 2β | 591.27 | 591.26 | 0.01 |

**BECC470s**
Chemical Formula: $C_{100}H_{186}N_2O_{22}P^-$
Exact Mass: 1798.32

**S1. Tandem Mass Spectrometry analysis of Synthetic BECC438s and BECC470s.** Synthetically synthesized Lipid A molecules were analyzed by tandem mass spectrometry to determine the structure.

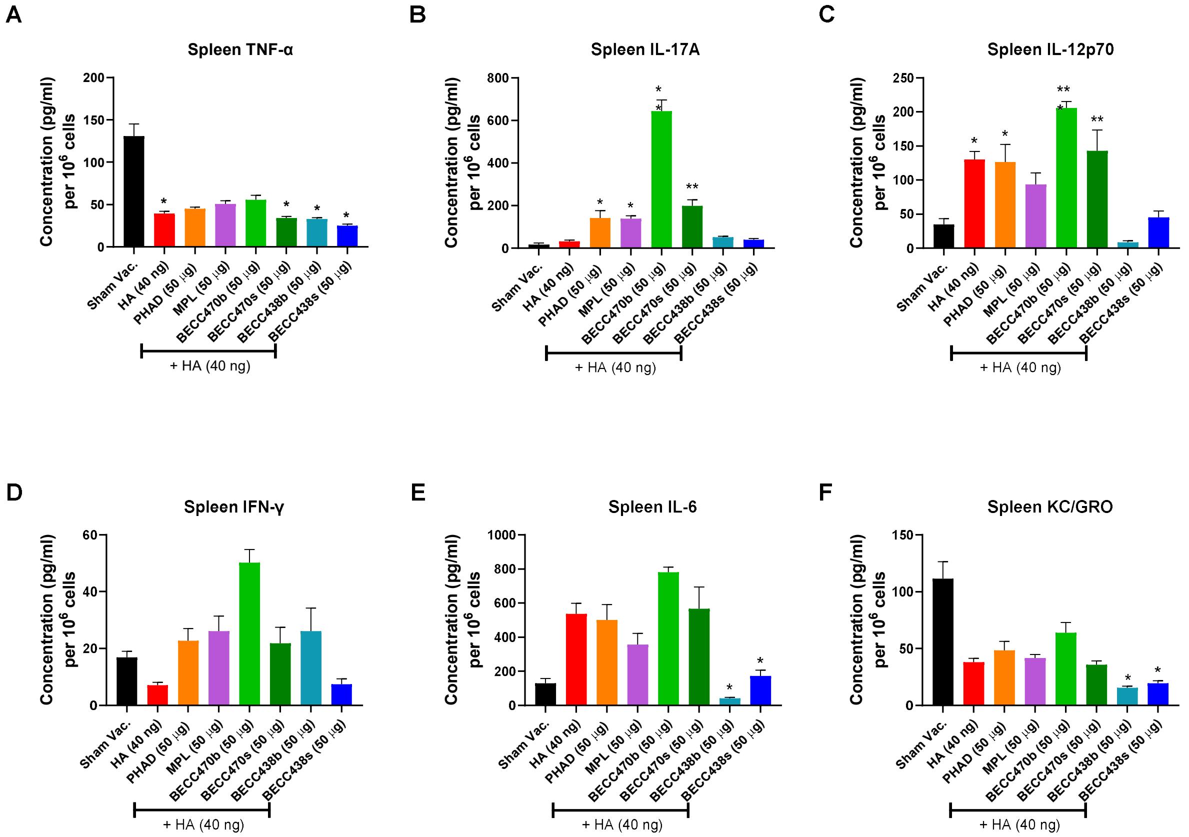

**S2. Cytokine induction by BECC adjuvanted HA pre-challenge.** 6-week-old BALB/c mice (n=5/group) were immunized on day 0 and day 14. On day 35 post immunization, mice were sacrificed, and single suspensions of spleen were prepared. Cell concentrations were equalized and were plated onto 96 well plates in the presence of HA. Supernatants were removed after 48 hours and analyzed for presence of cytokines (**A-F**). Data were analyzed by Dunnett's multiple comparison test by comparing to Sham Vaccinated/infected group. *p<0.05, **p<0.01, ***p<0.001.

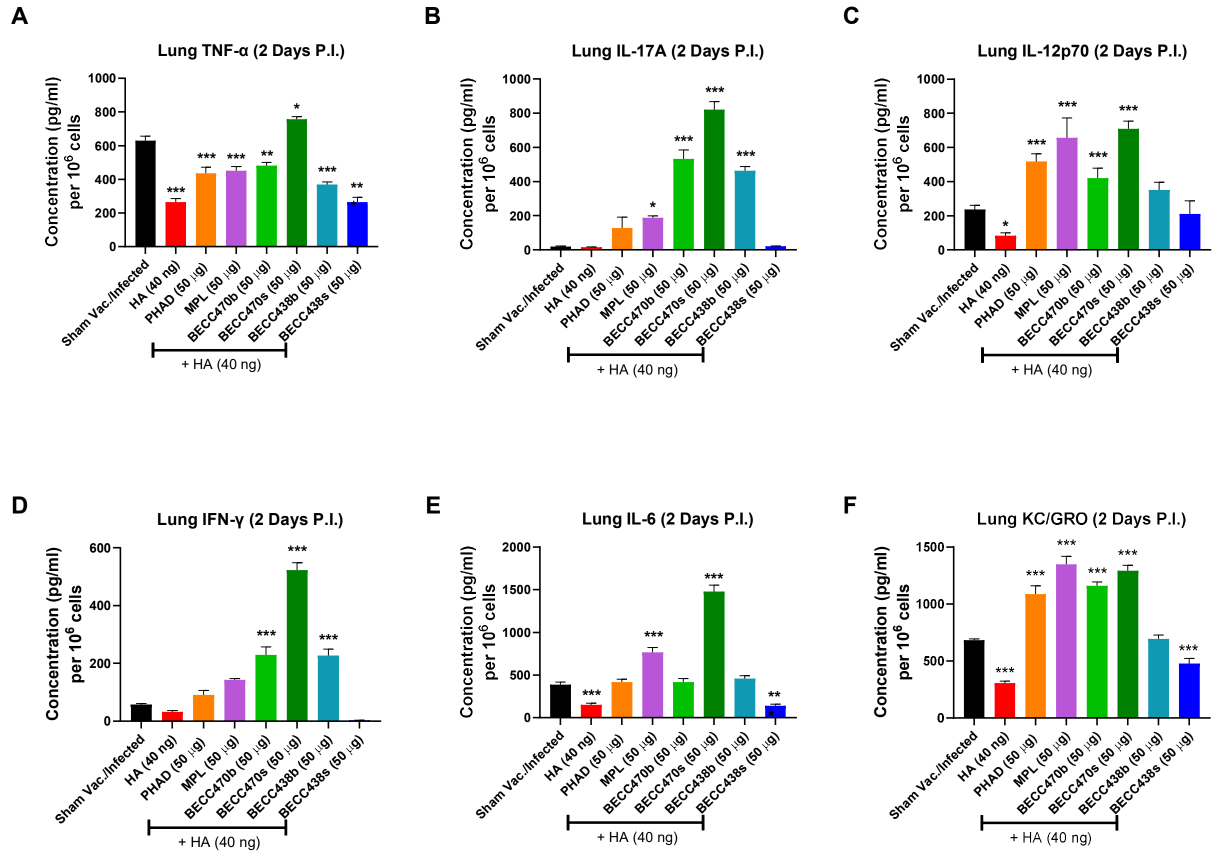

**S3. Cytokine induction by BECC adjuvanted HA post-challenge.** 6-week-old BALB/c mice (n=5/group) were immunized on day 0 and day 14. On day 35 post immunization, mice were intranasally challenged with 3200 PFU of NL09. Mice were sacrificed after 48 hours, and single suspensions of lungs were prepared. Cell concentrations were equalized and were plated onto 96 well plates in the presence of HA. Supernatants were removed after 48 hours and analyzed for presence of cytokines (**G-L**). Data were analyzed by Dunnett's multiple comparison test by comparing to Sham Vaccinated/infected group. *p<0.05, **p<0.01, ***p<0.001

| Table S1: List of Peptides (1/4) | | |
|---|---|---|
| **Peptide** | **Length** | **Sequence** |
| 1 of 139 | 15 | 1-MKAILVVLLYTFATA-15 |
| 2 of 139 | 15 | 5-LVVLLYTFATANADT-19 |
| 3 of 139 | 15 | 9-LYTFATANADTLCIG-23 |
| 4 of 139 | 15 | 13-ATANADTLCIGYHAN-27 |
| 5 of 139 | 15 | 17-ADTLCIGYHANNSTD-31 |
| 6 of 139 | 15 | 21-CIGYHANNSTDTVDT-35 |
| 7 of 139 | 15 | 25-HANNSTDTVDTVLEK-39 |
| 8 of 139 | 15 | 29-STDTVDTVLEKNVTV-43 |
| 9 of 139 | 15 | 33-VDTVLEKNVTVTHSV-47 |
| 10 of 139 | 15 | 37-LEKNVTVTHSVNLLE-51 |
| 11 of 139 | 15 | 41-VTVTHSVNLLEDKHN-55 |
| 12 of 139 | 15 | 45-HSVNLLEDKHNGKLC-59 |
| 13 of 139 | 15 | 49-LLEDKHNGKLCKLRG-63 |
| 14 of 139 | 15 | 53-KHNGKLCKLRGVAPL-67 |
| 15 of 139 | 15 | 57-KLCKLRGVAPLHLGK-71 |
| 16 of 139 | 15 | 61-LRGVAPLHLGKCNIA-75 |
| 17 of 139 | 15 | 65-APLHLGKCNIAGWIL-79 |
| 18 of 139 | 15 | 69-LGKCNIAGWILGNPE-83 |
| 19 of 139 | 15 | 73-NIAGWILGNPECESL-87 |
| 20 of 139 | 15 | 77-WILGNPECESLSTAS-91 |
| 21 of 139 | 15 | 81-NPECESLSTASSWSY-95 |
| 22 of 139 | 15 | 85-ESLSTASSWSYIVET-99 |
| 23 of 139 | 15 | 89-TASSWSYIVETPSSD-103 |
| 24 of 139 | 15 | 93-WSYIVETPSSDNGTC-107 |
| 25 of 139 | 15 | 97-VETPSSDNGTCYPGD-111 |
| 26 of 139 | 15 | 101-SSDNGTCYPGDFIDY-115 |
| 27 of 139 | 15 | 105-GTCYPGDFIDYEELR-119 |
| 28 of 139 | 15 | 109-PGDFIDYEELREQLS-123 |
| 29 of 139 | 15 | 113-IDYEELREQLSSVSS-127 |
| 30 of 139 | 15 | 117-ELREQLSSVSSFERF-131 |
| 31 of 139 | 15 | 121-QLSSVSSFERFEIFP-135 |
| 32 of 139 | 15 | 125-VSSFERFEIFPKTSS-139 |
| 33 of 139 | 15 | 129-ERFEIFPKTSSWPNH-143 |
| 34 of 139 | 15 | 133-IFPKTSSWPNHDSNK-147 |
| 35 of 139 | 15 | 137-TSSWPNHDSNKGVTA-151 |

| Table S1: List of Peptides (2/4) | | |
|---|---|---|
| **Peptide** | **Length** | **Sequence** |
| 36 of 139 | 15 | 141-PNHDSNKGVTAACPH-155 |
| 37 of 139 | 15 | 145-SNKGVTAACPHAGAK-159 |
| 38 of 139 | 15 | 149-VTAACPHAGAKSFYK-163 |
| 39 of 139 | 15 | 153-CPHAGAKSFYKNLIW-167 |
| 40 of 139 | 15 | 157-GAKSFYKNLIWLVKK-171 |
| 41 of 139 | 15 | 161-FYKNLIWLVKKGNSY-175 |
| 42 of 139 | 15 | 165-LIWLVKKGNSYPKLS-179 |
| 43 of 139 | 15 | 169-VKKGNSYPKLSKSYI-183 |
| 44 of 139 | 15 | 173-NSYPKLSKSYINDKG-187 |
| 45 of 139 | 15 | 177-KLSKSYINDKGKEVL-191 |
| 46 of 139 | 15 | 181-SYINDKGKEVLVLWG-195 |
| 47 of 139 | 15 | 185-DKGKEVLVLWGIHHP-199 |
| 48 of 139 | 15 | 189-EVLVLWGIHHPSTSA-203 |
| 49 of 139 | 15 | 193-LWGIHHPSTSADQQS-207 |
| 50 of 139 | 15 | 197-HHPSTSADQQSLYQN-211 |
| 51 of 139 | 15 | 201-TSADQQSLYQNADAY-215 |
| 52 of 139 | 15 | 205-QQSLYQNADAYVFVG-219 |
| 53 of 139 | 15 | 209-YQNADAYVFVGSSRY-223 |
| 54 of 139 | 15 | 213-DAYVFVGSSRYSKKF-227 |
| 55 of 139 | 15 | 217-FVGSSRYSKKFKPEI-231 |
| 56 of 139 | 15 | 221-SRYSKKFKPEIAIRP-235 |
| 57 of 139 | 15 | 225-KKFKPEIAIRPKVRD-239 |
| 58 of 139 | 15 | 229-PEIAIRPKVRDQEGR-243 |
| 59 of 139 | 15 | 233-IRPKVRDQEGRMNYY-247 |
| 60 of 139 | 15 | 237-VRDQEGRMNYYWTLV-251 |
| 61 of 139 | 15 | 241-EGRMNYYWTLVEPGD-255 |
| 62 of 139 | 15 | 245-NYYWTLVEPGDKITF-259 |
| 63 of 139 | 15 | 249-TLVEPGDKITFEATG-263 |
| 64 of 139 | 15 | 253-PGDKITFEATGNLVV-267 |
| 65 of 139 | 15 | 257-ITFEATGNLVVPRYA-271 |
| 66 of 139 | 15 | 261-ATGNLVVPRYAFAME-275 |
| 67 of 139 | 15 | 265-LVVPRYAFAMERNAG-279 |
| 68 of 139 | 15 | 269-RYAFAMERNAGSGII-283 |
| 69 of 139 | 15 | 273-AMERNAGSGIIISDT-287 |
| 70 of 139 | 15 | 277-NAGSGIIISDTPVHD-291 |
| 71 of 139 | 15 | 281-GIIISDTPVHDCNTT-295 |
| 72 of 139 | 15 | 285-SDTPVHDCNTTCQTP-299 |
| 73 of 139 | 15 | 289-VHDCNTTCQTPKGAI-303 |
| 74 of 139 | 15 | 293-NTTCQTPKGAINTSL-307 |
| 75 of 139 | 15 | 297-QTPKGAINTSLPFQN-311 |
| 76 of 139 | 15 | 301-GAINTSLPFQNIHPI-315 |
| 77 of 139 | 15 | 305-TSLPFQNIHPITIGK-319 |
| 78 of 139 | 15 | 309-FQNIHPITIGKCPKY-323 |
| 79 of 139 | 15 | 313-HPITIGKCPKYVKST-327 |
| 80 of 139 | 15 | 317-IGKCPKYVKSTKLRL-331 |

| Table S1: List of Peptides (3/4) | | |
|---|---|---|
| **Peptide** | **Length** | **Sequence** |
| 81 of 139 | 15 | 321-PKYVKSTKLRLATGL-335 |
| 82 of 139 | 15 | 325-KSTKLRLATGLRNIP-339 |
| 83 of 139 | 15 | 329-LRLATGLRNIPSIQS-343 |
| 84 of 139 | 15 | 333-TGLRNIPSIQSRGLF-347 |
| 85 of 139 | 15 | 337-NIPSIQSRGLFGAIA-351 |
| 86 of 139 | 15 | 341-IQSRGLFGAIAGFIE-355 |
| 87 of 139 | 15 | 345-GLFGAIAGFIEGGWT-359 |
| 88 of 139 | 15 | 349-AIAGFIEGGWTGMVD-363 |
| 89 of 139 | 15 | 353-FIEGGWTGMVDGWYG-367 |
| 90 of 139 | 15 | 357-GWTGMVDGWYGYHHQ-371 |
| 91 of 139 | 15 | 361-MVDGWYGYHHQNEQG-375 |
| 92 of 139 | 15 | 365-WYGYHHQNEQGSGYA-379 |
| 93 of 139 | 15 | 369-HHQNEQGSGYAADLK-383 |
| 94 of 139 | 15 | 373-EQGSGYAADLKSTQN-387 |
| 95 of 139 | 15 | 377-GYAADLKSTQNAIDE-391 |
| 96 of 139 | 15 | 381-DLKSTQNAIDEITNK-395 |
| 97 of 139 | 15 | 385-TQNAIDEITNKVNSV-399 |
| 98 of 139 | 15 | 389-IDEITNKVNSVIEKM-403 |
| 99 of 139 | 15 | 393-TNKVNSVIEKMNTQF-407 |
| 100 of 139 | 15 | 397-NSVIEKMNTQFTAVG-411 |
| 101 of 139 | 15 | 401-EKMNTQFTAVGKEFN-415 |
| 102 of 139 | 15 | 405-TQFTAVGKEFNHLEK-419 |
| 103 of 139 | 15 | 409-AVGKEFNHLEKRIEN-423 |
| 104 of 139 | 15 | 413-EFNHLEKRIENLNKK-427 |
| 105 of 139 | 15 | 417-LEKRIENLNKKVDDG-431 |
| 106 of 139 | 15 | 421-IENLNKKVDDGFLDI-435 |
| 107 of 139 | 15 | 425-NKKVDDGFLDIWTYN-439 |
| 108 of 139 | 15 | 429-DDGFLDIWTYNAELL-443 |
| 109 of 139 | 15 | 433-LDIWTYNAELLVLLE-447 |
| 110 of 139 | 15 | 437-TYNAELLVLLENERT-451 |
| 111 of 139 | 15 | 441-ELLVLLENERTLDYH-455 |
| 112 of 139 | 15 | 445-LLENERTLDYHDSNV-459 |
| 113 of 139 | 15 | 449-ERTLDYHDSNVKNLY-463 |
| 114 of 139 | 15 | 453-DYHDSNVKNLYEKVR-467 |
| 115 of 139 | 15 | 457-SNVKNLYEKVRSQLK-471 |
| 116 of 139 | 15 | 461-NLYEKVRSQLKNNAK-475 |
| 117 of 139 | 15 | 465-KVRSQLKNNAKEIGN-479 |
| 118 of 139 | 15 | 469-QLKNNAKEIGNGCFE-483 |
| 119 of 139 | 15 | 473-NAKEIGNGCFEFYHK-487 |
| 120 of 139 | 15 | 477-IGNGCFEFYHKCDNT-491 |
| 121 of 139 | 15 | 481-CFEFYHKCDNTCMES-495 |
| 122 of 139 | 15 | 485-YHKCDNTCMESVKNG-499 |
| 123 of 139 | 15 | 489-DNTCMESVKNGTYDY-503 |
| 124 of 139 | 15 | 493-MESVKNGTYDYPKYS-507 |
| 125 of 139 | 15 | 497-KNGTYDYPKYSEEAK-511 |

| Table S1: List of Peptides (4/4) | | |
|---|---|---|
| **Peptide** | **Length** | **Sequence** |
| 126 of 139 | 15 | 501-YDYPKYSEEAKLNRE-515 |
| 127 of 139 | 15 | 505-KYSEEAKLNREEIDG-519 |
| 128 of 139 | 15 | 509-EAKLNREEIDGVKLE-523 |
| 129 of 139 | 15 | 513-NREEIDGVKLESTRI-527 |
| 130 of 139 | 15 | 517-IDGVKLESTRIYQIL-531 |
| 131 of 139 | 15 | 521-KLESTRIYQILAIYS-535 |
| 132 of 139 | 15 | 525-TRIYQILAIYSTVAS-539 |
| 133 of 139 | 15 | 529-QILAIYSTVASSLVL-543 |
| 134 of 139 | 15 | 533-IYSTVASSLVLVVSL-547 |
| 135 of 139 | 15 | 537-VASSLVLVVSLGAIS-551 |
| 136 of 139 | 15 | 541-LVLVVSLGAISFWMC-555 |
| 137 of 139 | 15 | 545-VSLGAISFWMCSNGS-559 |
| 138 of 139 | 15 | 549-AISFWMCSNGSLQCR-563 |
| 139 of 139 | 14 | 553-WMCSNGSLQCRICI-566 |
